# Orphan GPR84 facilitates uropod de-adhesion to terminate leukocyte diapedesis during inflammation

**DOI:** 10.64898/2026.01.28.702287

**Authors:** Clare Latta, Terrence M. Trinca, Francesca Robertson, Rachel Lau, Anna Barkaway, Loic Rolas, Matthew Golding, Paul R.C. Imbert, Haitao Wang, Almke Bader, Liam S. Hill, Lorna Hodgson, Barbara Walzog, Myles Lewis, Mathieu Benoit-Voisin, Paul Martin, Sussan Nourshargh, Helen Weavers

## Abstract

Leukocyte migration through venular walls is an essential component of effective immunity. While the key molecular players driving the initial steps of this response are known, the terminating signals remain unclear. Here, we have identified a conserved role for GPR84 family GPCRs in successful completion of the final stages of leukocyte extravasation through acutely inflamed vessels. The integration of high resolution intravital imaging with cell-specific genetics revealed that genetic deficiency of GPR84 orthologues in mice and *Drosophila*, results in defective detachment of transmigrating immune cells from vessel walls. Mechanistically, transcriptomics revealed GPR84-deficient neutrophils exhibit defective actin cytoskeletal regulation and cellular adhesion/de-adhesion. Consistent with this, our fly-murine pipeline shows that GPR84 supports localized and dynamic Rho activation to enable detachment of the immune cell uropod from vessel exit sites. Moreover, pharmacological blockade of GPR84 signaling dampened immune cell migration in multiple murine acute inflammatory settings. Collectively, our findings present GPR84 as a novel physiological regulator of immune cell extravasation that is amenable to therapeutic targeting for modulating leukocyte infiltration into inflamed tissues.

**Summary:** Here we identify the GPR84 family of GPCRs as key regulators of effective immune cell extravasation *in vivo*. Mechanistically, through integrating genetically tractable *Drosophila* and murine *in vivo* models, we show how leukocyte GPR84 supports dynamic Rho-dependent detachment of stretched uropods as these cells exit vessels.

## Introduction

A key rate-limiting step in the development of an acute inflammatory reaction is the emigration of innate immune cells (neutrophils and monocytes) from the blood circulation to sites of tissue infection or damage (Nourshargh and Alon, 2014; Muller, 2016; Vestweber, 2015). This response begins with the capture and adhesion of leukocytes to venular walls, as described by the classical adhesion cascade, with the final step being the breaching of vessel walls (commonly post-capillary venules) (Ley et al., 2007). The latter, termed immune cell extravasation (also known as transmigration or diapedesis) (Nourshargh and Alon, 2014; Nourshargh et al., 2010), is a dynamic process that involves the active participation of both immune and vascular cells (Nourshargh and Alon, 2014; Muller, 2016; Vestweber, 2015).

Murine genetic approaches, combined with high resolution intravital microscopy and complementary *in vitro* flow studies, have laid out the key steps and molecular players that underpin the initiation and progression of leukocyte extravasation (Muller, 2016; Vestweber, 2015; Nourshargh and Alon, 2014; Ley et al., 2007; Phillipson et al., 2006). To exit the blood circulation, immune cells are required to breach different components of the venular wall, namely endothelial cells (ECs) that line the inner aspect of all blood vessels, as well as pericytes and the venular basement membrane (BM). Characterization of immune cell and EC/pericyte expressed adhesion molecules, most notably proteins expressed at EC junctions (such as PECAM-1, JAM-A, JAM-C and VE-cadherin), and immobilized chemokines, have shed much light on the underlying molecular mechanisms (Vestweber, 2015; Nourshargh and Alon, 2014; Muller, 2016; Ley et al., 2007). In breaching the endothelial layer, leukocytes commonly migrate through EC junctions (paracellular), and less frequently in the peripheral circulation, they exhibit migration through the body of individual ECs (transcellular). Once in the sub-EC space, leukocytes probe for weak spots in the basement membrane and gaps in the pericyte sheath, as directed by chemotactic cues from vascular and perivascular cells before fully exiting the venular wall and commencing their extravascular migration (Joulia et al., 2022; Girbl et al., 2018; Proebstl et al., 2012).

Despite our in-depth understanding of this process, the signaling events that enable successful completion of each sequential step of leukocyte extravasation remain unclear. Unravelling such dynamic molecular cascades *in vivo* at the single cell level necessitate tractable models amenable to high resolution live imaging and cell-specific genetic manipulation. Whilst *Drosophila* were not previously considered an appropriate model to investigate immune cell extravasation because of their open circulation (Razzell et al., 2011), we recently identified a period during their pupal development when the beating wing hearts pulse blood (including the hemocytes, *Drosophila* leukocytes) through the developing wing vessels (known as wing ‘veins’) (Thuma et al., 2018). Here, wounding of the adjacent epithelium triggers the recruitment of hemocytes across the vessel wall that can be imaged in real-time with high spatiotemporal resolution (Thuma et al., 2018). Capitalizing on this model, and harnessing *Drosophila’s* unrivalled genetic tractability, we identified *Tre1* (*Trapped in endoderm 1*), a Rhodopsin family G protein–coupled receptor (GPCR), as essential for hemocyte extravasation across vessels walls. Additionally, live imaging studies revealed a failure of Tre1-deficient hemocytes to fully extravasate, suggesting a non-redundant role for Tre1 in modulating immune cell extravasation machinery (Thuma et al., 2018).

In the present study we have undertaken an in-depth characterization of the functions of GPR84, the closest mammalian orthologue of Tre1, in leukocyte extravasation in murine models of inflammation using both genetic and pharmacological strategies. GPR84, an orphan GPCR of the rhodopsin-like Class A family (Wittenberger et al., 2001), exhibits mRNA expression largely restricted to peripheral immune cells (predominantly myeloid cells and microglia in the CNS) and is upregulated in certain inflammatory settings (Bouchard et al., 2007; Recio et al., 2018; Yousefi et al., 2001). While GPR84 has been identified as a pro-inflammatory immune cell receptor in experimental models of inflammation (Recio et al., 2018; Cooper et al., 2024; Puengel et al., 2020; Abdel-Aziz et al., 2016; Gagnon et al., 2018; Wang et al., 2023a), its precise role in mediating key steps in immune cell migration at the single cell level is unknown.

To address this, we have integrated genetic, live imaging, transcriptomic, electron microscopy and pharmacological studies to shed light on the subcellular mechanisms through which GPR84 orthologues exhibit conserved functions during leukocyte extravasation. Briefly, genetic knockout of murine GPR84 impaired all stages of neutrophil diapedesis, from breaching the endothelium and the pericyte layer through to final detachment from the venular wall. Strikingly, live imaging and correlative light electron microscopy (CLEM) revealed a key impairment in GPR84 deficient leukocytes during the final stages of extravasation, namely failure of the leukocyte uropod to detach from the vessel wall. Our mechanistic studies involving both *Drosophila* and murine models suggest that Tre1/GPR84 plays an essential role in promoting Rho GTPase-driven release of integrin-mediated adhesions. Consistent with this, pharmacological blockade of GPR84 signaling *in vivo* significantly dampened neutrophil recruitment in multiple acute inflammatory reactions, including experimental models of cutaneous ischemia-reperfusion injury, peritonitis and skin wound healing. Collectively our observations identify a distinct role for the conserved GPCR84 family as rate limiting players in immune cell extravasation, with potential for therapeutic targeting for pathologies characterized by overexuberant inflammation.

## Results

### The GPCR Tre1 is required for effective hemocyte extravasation from *Drosophila* pupal veins

We recently demonstrated that *Drosophila* pupal wings in which hemocytes circulate through vessels (the wing ‘veins’), offer the opportunity to model immune cell extravasation in response to an inflammatory stimulus in the form of a small epithelial wound adjacent to a wing vein (**Figure 1A-D**) (Thuma et al., 2018). Within this model, we can observe hemocytes circulating within the hemolymph pulsing through wing vessels, with vessel LV3 (that extends in one focal plane) being particularly amenable for imaging (**Figure 1B-C**). Within 10 min of making a laser wound in the epithelium adjacent to this vessel, hemocytes are seen to probe the vessel wall in the vicinity of the injury before extravasating and subsequently migrating towards the wound (**Figure 1D**).

**Figure 1.**
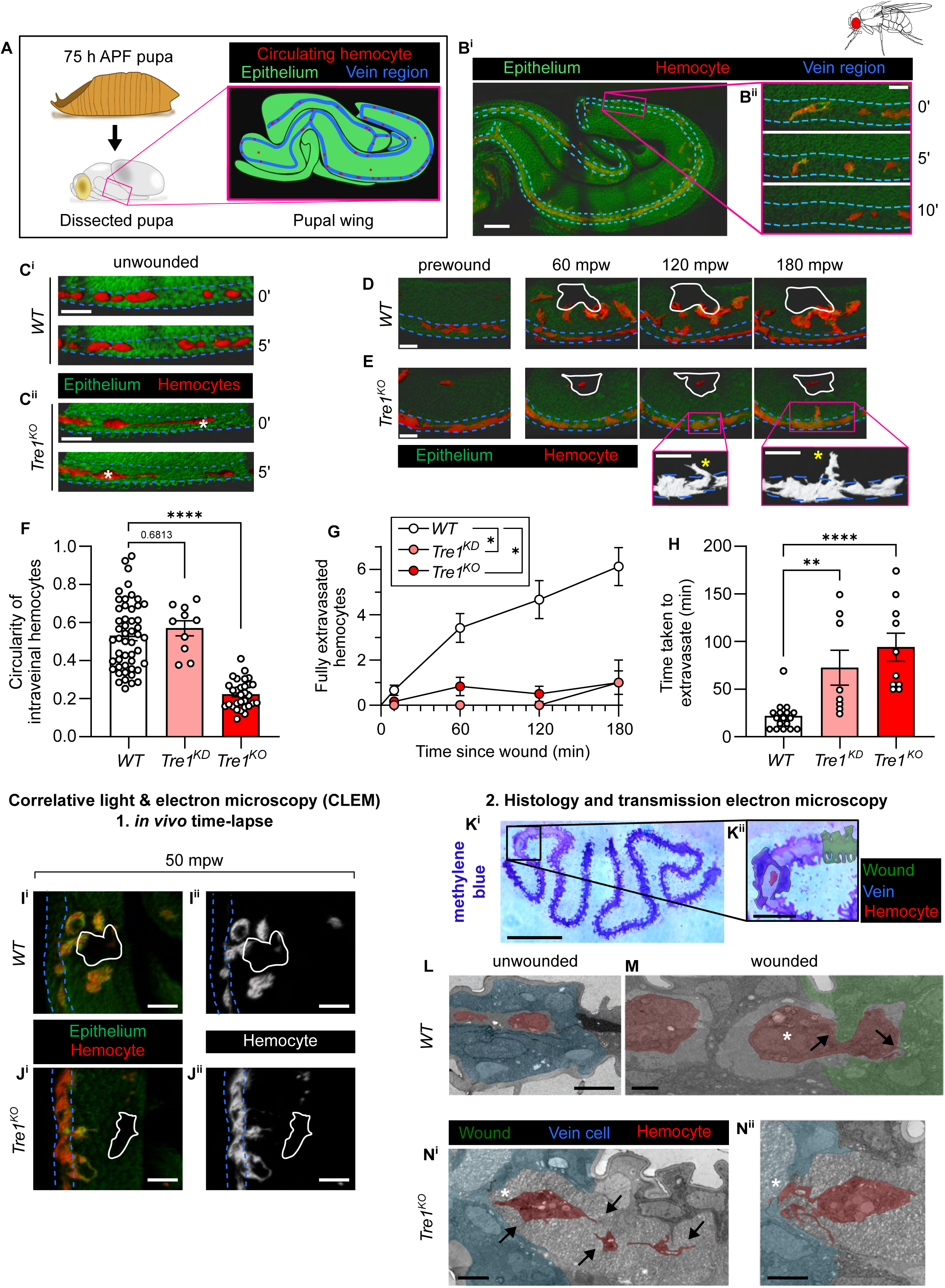
*Drosophila* late-pupal model of immune cell extravasation. **(A)** Schematic to illustrate how 75h APF (after puparium formation) pupae are dissected to enable confocal imaging of the developing wings. **(B^i^)** Representative *in vivo* imaging of whole-pupal wing with epithelium labelled in green (*ubi-GMA*), hemocytes labelled in red (*srp-Gal4>mCherry; srp-mch*), and vein regions indicated with a blue dotted line. **(B^ii^)** Time-lapse IMARIS renderings of a vein region with circulating hemocytes. **(C^i^ and C^ii^)** Comparison of *Oregon R* (*WT*) and *Tre1^KO^* hemocytes’ morphology as they circulate in unwounded conditions. *Tre1^KO^*hemocytes have cytoplasmic extensions that appear stuck to the luminal side of the vein cells (white asterisks). **(D)** Timelapse confocal images of *WT* hemocytes (red) responding to a laser wound (solid white outline). **(E)** By contrast, *Tre1^KO^* hemocytes cannot fully dissociate from the vessel wall. High magnification renderings (from regions in magenta boxes) illustrate the distinct morphology of mutant hemocytes and their long leading edges (yellow asterisks). **(F)** Quantification of the circularity of intravenous circulating *WT*, *Tre1^KD^* and *Tre1^KO^*hemocytes; bar plots represent calculated mean for each genotype, and dots represent individual hemocytes pooled from multiple animals. **(G)** Quantification of numbers of hemocytes that have fully dissociated from veins and extravasated towards wounds. **(H)** Analysis of time taken for individual hemocytes to extravasate from veins; bar plots represent calculated mean for each genotype, and dots represent individual hemocytes pooled from multiple animals. **(I - J)** *In vivo* confocal imaging of *WT* **(I and I^i^)** versus *Tre1^KO^* **(J and J^i^)** hemocytes as they extravasate towards wounds (white outline), just prior to fixation at 50 minutes post wounding (mpw) for further EM analysis in a correlative light and electron microscopic (CLEM) study. **(K^i^)** Methylene blue stained resin-embedded transverse sections of a *WT* wounded sample. **(K^ii^)** Higher magnification view (from black boxed region) reveals a single hemocyte (false coloured red) within the lumen of the vein (blue) with a wound (green) adjacent to the vessel. **(L)** Transmission electron microscopy (TEM) of an unwounded *WT* vein (blue) with two hemocytes (red) within. **(M)** Wounded *WT* hemocytes (red) 15 minutes *post* extravasation as they traverse the intervein epithelium towards a wound (green). **(N^i^ and N^ii^)** *Tre1^KO^* hemocytes have partially extravasated and exhibit extensive leading edges (black arrows), but remain adherent to the vein cells, with multiple lagging tails observed (white asterisks). Data in plots (**F-H**) were analyzed using one-way ANOVA with a Dunnett’s test for multiple comparisons. For each quantification data represents at least 9 animals per any genotype and represented the mean ± SEM. Asterisks = *P<0.05, **P<0.01, ***P<0.001, ****P<0.0001. Scale bars: B and K^i^ = 50 µm; B^i^, C^i^, C^ii^, D, E, I^i^ - J^ii^, K^ii^ = 15 µm; L = 5 µm; M, N^i^ and N^ii^ = 2 µm. Also see **Supplementary Movie S1.**

Harnessing the live imaging opportunities of the translucent pupa, together with *Drosophila’s* genetic tractability, we sought novel genes that might drive immune cell extravasation. Here, extending our previous works (Thuma et al., 2018), and utilizing both a hemocyte-specific Tre1-RNAi knockdown (Tre1^KD^) approach and a Tre1 null (Tre1^KO^) mutant (in which the endogenous Tre1 locus has been removed; **Supplementary Figure 1A)** (Kim et al., 2021), our live imaging studies show Tre1 to be fundamental for complete hemocyte extravasation (**Figure 1C-E**). Within the wing veins, hemocytes lacking Tre1 showed an intravenous morphological phenotype even in unwounded conditions, exhibiting cytoplasmic extensions that appeared stuck to the luminal side of the vein cells resulting in reduced circularity **(Figure 1C and F)**, a sustained phenomenon that was evident for the duration of the imaging period.

Following wounding proximal to the wing vein structure, hemocytes in Tre1 null pupae showed an extreme phenotype, with fewer hemocytes fully extravasating and those that did fully extravasate, taking longer when compared to the responses detected in the controls or RNAi-based knockdown experiments (**Figure 1D-H, Supplementary Figure 1B-C** and **Supplementary Movie S1**). The tails of Tre1-deficient hemocytes remained adherent to the vein walls, suggesting that Tre1 may specifically regulate the perivascular detachment phase of hemocyte extravasation (**Figure 1E**). We next undertook correlative light and transmission electron microscope imaging (CLEM) studies to compare hemocyte morphologies post wounding in control and Tre1-deficient pupae (**Figure 1I-N**). Timelapse movies enabled the capture of hemocytes in the act of extravasating *in vivo* (**Figure 1I-J**) and these samples were immediately prepared to visualize this process at the ultrastructural level via transmission electron microscopy (**Figure 1K-N**). In control unwounded pupae, we observed hemocytes within the lumen of the wing vein (**Figure 1L**); following wounding, hemocytes were observed outside of the vessel exhibiting relatively short lamellipodia en route to the wound (**Figure 1M**). Tre1-deficient hemocytes, however, showed long leading extensions and retained multiple lagging tethers linking them to the vein wall cells and to the interior of the vein (**Figure 1N^i^ and N^ii^).** Consistent with our live-imaging studies, these CLEM data suggest that hemocytes are disrupted in their capacity to extravasate completely through wing vessels after a sterile injury, potentially due to an incapacity to de-adhere from vessel cells.

### Tre1 shares structural homology to its conserved mammalian orthologue, GPR84

To assess whether our *Drosophila* model could identify extravasation players with conserved roles in mammals, we explored whether the human and mouse genomes possess Tre1 orthologues. Phylogenetic analysis of amino acid sequences revealed several *Drosophila* paralogues and mammalian orthologues of Tre1 (**Figure 2A**), including GPR84 with 34% amino acid sequence homology. Neither Tre1, nor its mammalian orthologues, have experimentally derived structures, but AlphaFold enabled us to compare their predicted structures (**Figure 2B and Supplementary Movie S2**) (Jumper et al., 2021). This *in silico* comparison suggested high structural homology with significant identical elements (yellow, **Figure 2B**), suggesting likely shared functionality.

**Figure 2.**
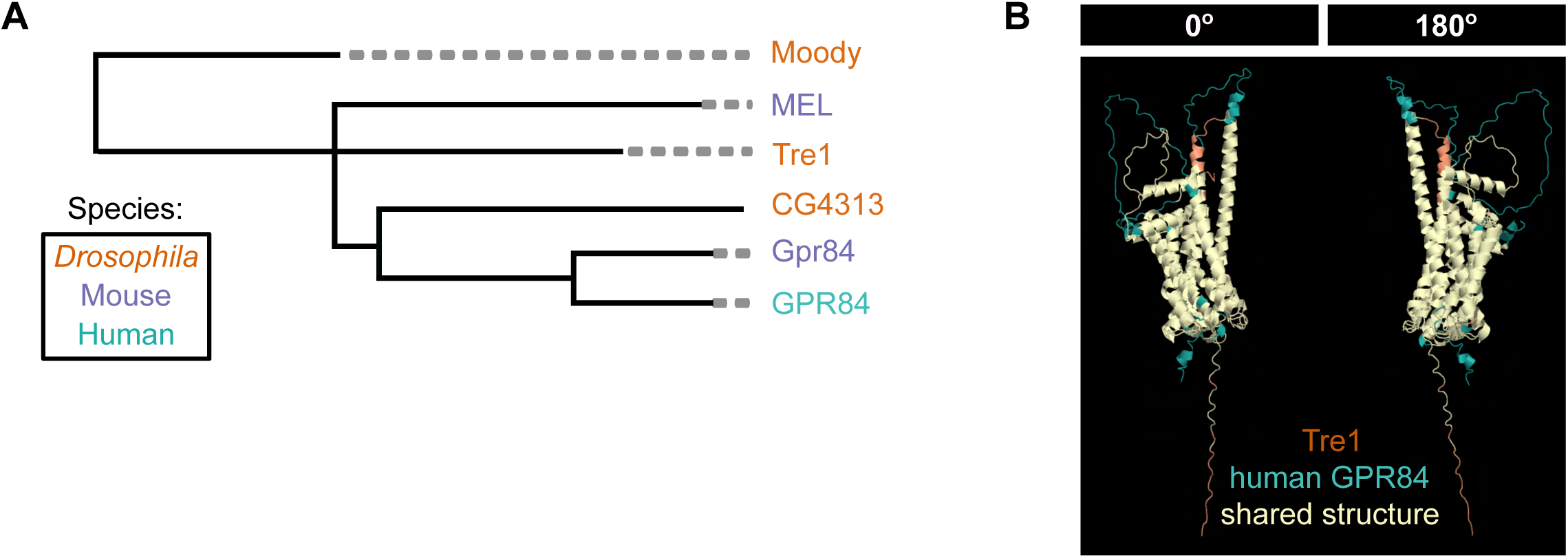
Tre1 is related to and shares structural homology with its mammalian orthologue GPR84. **(A)** Phylogenetic tree constructed from multiple sequence alignment between predicted Tre1 paralogues in *Drosophila* (maroon), and orthologues in mice (blue) and humans (teal). The size of horizontal branches corresponds to the estimated level of evolutionary conservation between orthologues on adjacent nodes. **(B)** Structural comparison of AlphaFold3 predicted models (two orientations: back and front) for Tre1 (maroon) and human GPR84 (teal), with their extensively shared homologous structural regions indicated (yellow). Structures were aligned with UCSF ChimeraX Matchmaker (Pettersen et al., 2021); the alignment yielded 238 Cα pairs with a RMSD of 1.14 Å (1.135 Å, pruned) and across all 314 aligned pairs, the RMSD was 4.10 Å. Also see **Supplementary Movie S2.**

### GPR84 mediates neutrophil extravasation in a murine model of acute inflammation

Extending the *Drosophila* works, we sought to investigate the functional role of the predicted mammalian orthologue of Tre1, GPR84. In initial studies, aiming to unravel the stage of immune cell migration mediated by GPR84, we analysed immune cell behaviour of WT and *Gpr84^-/-^* mice in control or acutely-inflamed mouse cremaster muscles by brightfield intravital microscopy (IVM) (**Figure 3A-D** and **Supplementary Movie S3**). Here, whilst local TNF (4 h) induced a significant and comparable level of intraluminal leukocyte adhesion in both WT and *Gpr84^-/-^*mice (**Figure 3C**), TNF-induced leukocyte extravasation was significantly reduced in *Gpr84^-/-^*mice as compared to WT littermates **(Figure 3D**). Furthermore, chimeric animals with selective myeloid GPR84 deficiency (generated through transfer of bone marrow cells to irradiated recipients; **Figure 3E**), showed defective neutrophil extravasation (**Figure 3F-G**). Of note, we saw no difference in the total or differential leukocyte counts in circulating blood of WT and KO mice under basal conditions (**Supplementary Figure 2A**) and the chimeric mouse lines showed equal reconstitution efficiency (**Supplementary Figure 2B-C**). Collectively, the present findings illustrate a specific and cell-autonomous role for GPR84 in regulating the passage of leukocytes through inflamed venular walls.

**Figure 3.**
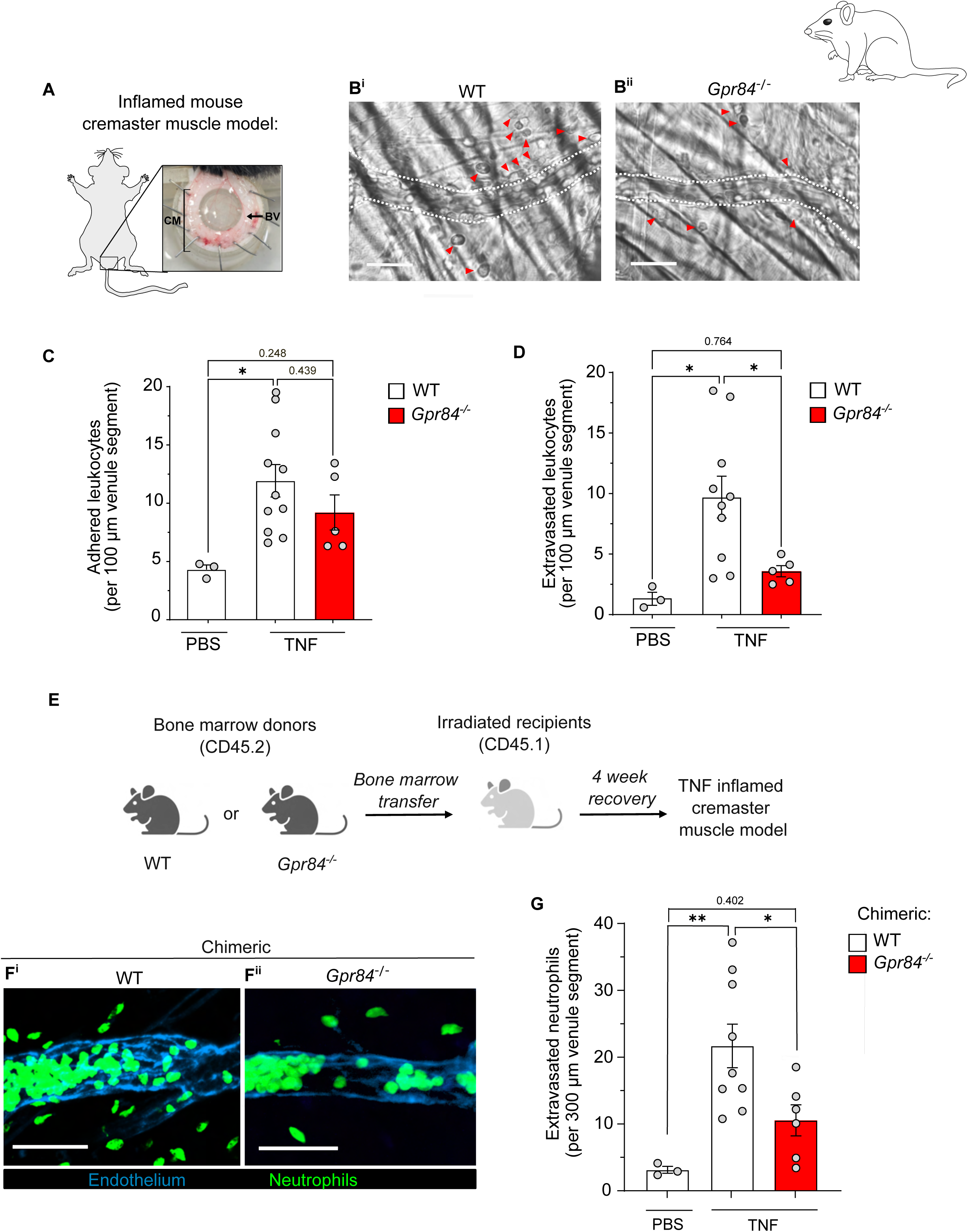
Myeloid cell GPR84-deficiency reduces neutrophil extravasation in TNF-stimulated cremaster muscle. **(A)** Schematic shows the exteriorised murine cremaster muscle (CM), subjected to TNF-induced inflammation immediately prior to intravital microscopy analysis. Arrow indicates the cremasteric microvasculature and presence of blood vessels (BV). **(B)** Representative images of **(B^i^)** WT and **(B^ii^)** *Gpr84^-/-^* cremasteric post capillary venules subjected to intrascrotal injection of TNF and subsequently imaged by brightfield intravital microscopy (IVM) at 4 h. Arrows indicate the presence of extravasated leukocytes within the interstitial tissue. **(C and D)** Number of firmly adhered **(C)** and extravasated **(D)** leukocytes in WT and *Gpr84^-/-^* mice (n = 3 – 11 mice/group); each data point represents the average number of leukocytes from 2-8 venules per mouse. **(E)** Schematic depicting the generation of WT and *Gpr84*^-/-^ bone-marrow chimeric mice. **(F)** Inflamed cremaster muscles were fixed and immunostained for endothelial cells (PECAM-1; blue) and neutrophils (MRP-14; green). Representative confocal images of cremasteric postcapillary venules from **(F^i^)** WT and **(F^ii^)** *Gpr84^-/-^* chimeric mice. **(G)** Quantification of the average number of extravasated neutrophils from 8-13 venule segments in TNF-inflamed cremaster muscles of chimeric WT and *Gpr84*^-/-^ mice (n= 3-8 mice/group). Each data point represents an individual mouse. Data in C, D and G are Mean ± SEM and analysed by One-way ANOVA with a Tukey’s test for multiple comparisons. *P <0.05, **P < 0.01. Scale bars: B = 20 µm, F = 50 µm. Also see **Supplementary Movie S3**.

### GPR84-deficient neutrophils display excessive adhesive interactions within inflamed vessels

Having identified defective immune cell extravasation in both Tre1-deficient *Drosophila* and GPR84-deficient mice, we next sought to gain more insight into the mechanism of action of GPR84 in real-time and at the single cell level using high resolution 4D confocal IVM. For this purpose, we interbred *Gpr84^- /-^* mice with the compound mouse reporter strain *Lyz2-EGFP-ki*; *Acta2-RFPcherry-Tg* that exhibits endogenous GFP^high^ neutrophils and RFP^+^ pericytes/smooth muscle cells. For imaging ECs, or more specifically EC junctions, the mice received a local injection of non-blocking fluorescently labelled anti-PECAM-1 mAb, as previously described (Woodfin et al., 2011). The application of this platform to exteriorised TNF-stimulated cremaster muscles revealed aberrant neutrophil behaviours in GPR84 KO mice at multiple stages of neutrophil extravasation, all reminiscent of defective de-adhesion. Observation of initial responses (**Figure 4A**) revealed abnormal intraluminal neutrophil structures in GPR84 deficient mice, as compared to WT controls. This was most pronounced during the luminal crawling and adhesion phases (**Figure 4A-B**) where *Gpr84^-/-^* neutrophils displayed exaggerated tethering to the endothelium (**Figure 4B^i^-B^iii^).** These observations were similar to the aberrant intraluminal structures detected in Tre1 null *Drosophila* hemocytes (**Figure 1C and F**). Interestingly, neutrophil fragments were frequently detected in close apposition to sites of neutrophil adhesion in KO mice, a response that may allude to excessive neutrophil adhesion and difficulties with detachment from the endothelium **(Figure 4B^iii^)**.

**Figure 4.**
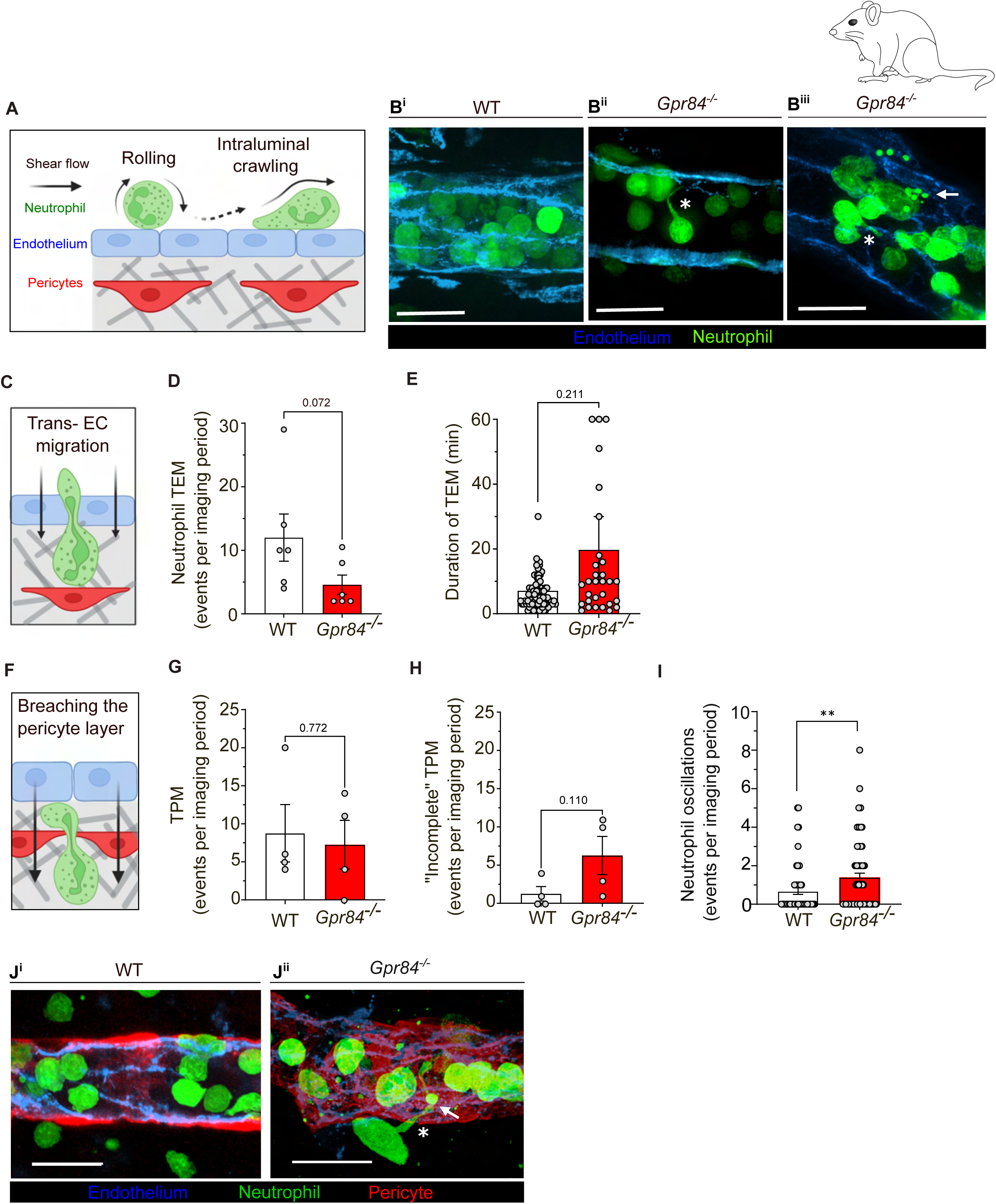
GPR84-deficient neutrophils display impaired trans-endothelial migration (TEM) and aberrant breaching of the pericyte layer during extravasation. (A-J) The cremaster microvasculature of WT and *Gpr84^-/-^* mice on a *Lyz2-EGFP-ki; Acta2-RFPcherry-Tg* genetic background were acutely inflamed with intrascrotal injection of TNF (300 ng) and subsequent extravasation events recorded by confocal IVM. **(A)** Schematic depicting the responses being analysed in Panel B, i.e. intraluminal neutrophil rolling and crawling. **(B)** Confocal images of luminal neutrophils in venules of WT **(B^i^)** and *Gpr84^-/-^* mice **(B^ii^-B^iii^)** 4 h post administration of TNF. **(B^ii^- B^iii^)** Asterisks indicate the presence of elongated neutrophil uropod structures tethered to endothelial cells and arrows show the increased deposition of neutrophil fragment structures in *Gpr84^-/-^* mice. **(C)** Schematic depicting the response being analysed in Panels D and E, i.e. trans endothelial cell migration (TEM). **(D-E)** Quantification of **(D)** the number of neutrophil TEM events within an imaging period of 2-5 h, each data point represents the average from 1-2 venules per mouse, and **(E)** duration of neutrophil TEM (in min), (n= 6-7 mice/group). Bar plots represent calculated mean value for each genotype, and data points represent single neutrophils pooled from multiple animals. **(F)** Schematic depicting the response being analysed in Panels G-I, i.e. trans-pericyte migration (TPM). **(G-I)** Number of neutrophils exhibiting TPM events **(G)** and (**H)** the number of incomplete TPM events, presented as the average from 1-2 venules per mouse within an imaging period, WT (n=4) and *Gpr84^-/-^* (n=4). **(I)** Number of neutrophil oscillations at the pericyte layer, each data point represents an individual neutrophil from 1-2 venules per mouse, WT (n=4) and *Gpr84^-/-^* (n=4). **(J)** Confocal images of neutrophil morphology when breaching the pericyte layer in **(J^i^)** WT and (J^ii^) *Gpr84^-/-^* mice. Asterisk indicates sub-endothelial neutrophil structures with **(J^ii^)** displaying an elongated tail retained within the sub-EC space and increased deposition of neutrophil fragmented structures (arrow). Data shows Mean ± SEM. Data was analysed by unpaired T-test (on the means per mouse) with significance shown by * P< 0.05, ** P< 0.01. Scale bars: B and J = 20 µm. (A, C and F) were created in BioRender. L, C. (2025) https://BioRender.com/bup6fba.

In analysing the frequency and dynamics of neutrophil trans-endothelial cell migration (TEM) (**Figure 4C)**, GPR84-deficient neutrophils consistently displayed a reduced number of TEM events (**Figure 4D**) and an extended duration to fully breach the endothelium (**Figure 4E**). Post breaching of the endothelium, KO mice subsequently showed multiple aberrant interactions between neutrophils and pericytes (**Figure 4F**). Specifically, despite no overall differences in the number of neutrophils breaching the pericyte layer (termed trans-pericyte migration; TPM) in WT and KO mice **(Figure 4G),** we noted an increase in the number of neutrophils that could not fully complete their migration across the pericyte layer **(Figure 4H).** Furthermore, KO neutrophils exhibited a distinct hesitancy and increased oscillatory behaviour, characterised by a repetitive probing and retracting of the neutrophil cell body, prior to eventual completion of their migration across the pericyte sheath (**Figure 4I**). In many instances, this wavering neutrophil-pericyte interaction was observed throughout the whole duration of the imaging period in GPR84 KO mice and this step ultimately failed (see **Supplementary Movie S4**). Interestingly, similar to the defective intraluminal responses, post breaching the pericyte layer, a subset of *Gpr84^-/-^*neutrophils again presented elongated tail-like structures within the sub-EC space (**Figure 4J**). The latter were commonly associated with evidence of neutrophil fragment deposition, supporting the notion of a generally impaired leukocyte de-adhesion within inflamed vessel walls of KO mice.

### GPR84-deficient neutrophils exhibit defective detachment from inflamed vessels

Guided by the striking phenotype of elongated tails observed in Tre1-deficient hemocytes (**Figure 1**), we extended our murine confocal IVM studies to analyse neutrophil detachment from inflamed venules in the perivascular space (**Figure 5A**). As a quantitative assessment of neutrophil morphology, we measured the maximum length of perivascular neutrophils during the imaging period (**Figure 5B**). This analysis revealed that in GPR84-deficient mice, perivascular neutrophils present two principal profiles, (i) a population exhibiting an ameboid morphology that shows almost total perivascular arrest, and (ii) a population with an extended and persistent uropod formation, reaching > 50 µm in length in some cells. The resultant effect was a delayed detachment of KO neutrophils from inflamed venular walls. Hence, in entering the perivascular space, while WT neutrophils displayed brief attachments (**∼**12 mins) to the exit site of the venular wall, in *Gpr84^-/-^* mice, **∼**80% of the perivascular neutrophils required > 30 minutes to fully detach and enter the surrounding interstitial space (**Figure 5C and Supplementary Movie S5)**. The aberrant GPR84 KO neutrophil tails **(Figure 5D)** were probed further at the ultrastructural level by transmission electron microscopy **(Figure 5E-F).** Whereas, in WT mice we saw fully detached neutrophils with short uropods (as if recently detached from adjacent pericytes) (**Figure 5E**), in *Gpr84^-/-^*mice we observed extensive lagging tails that appeared tethered to the pericyte wall **(Figure 5F^i^).** The latter were often associated with fibrillar matrix **(Figure 5F^ii^ - F^ii^)**. Collectively, the findings suggest exaggerated and prolonged leukocyte adhesion at multiple sites within and along the venular wall.

**Figure 5.**
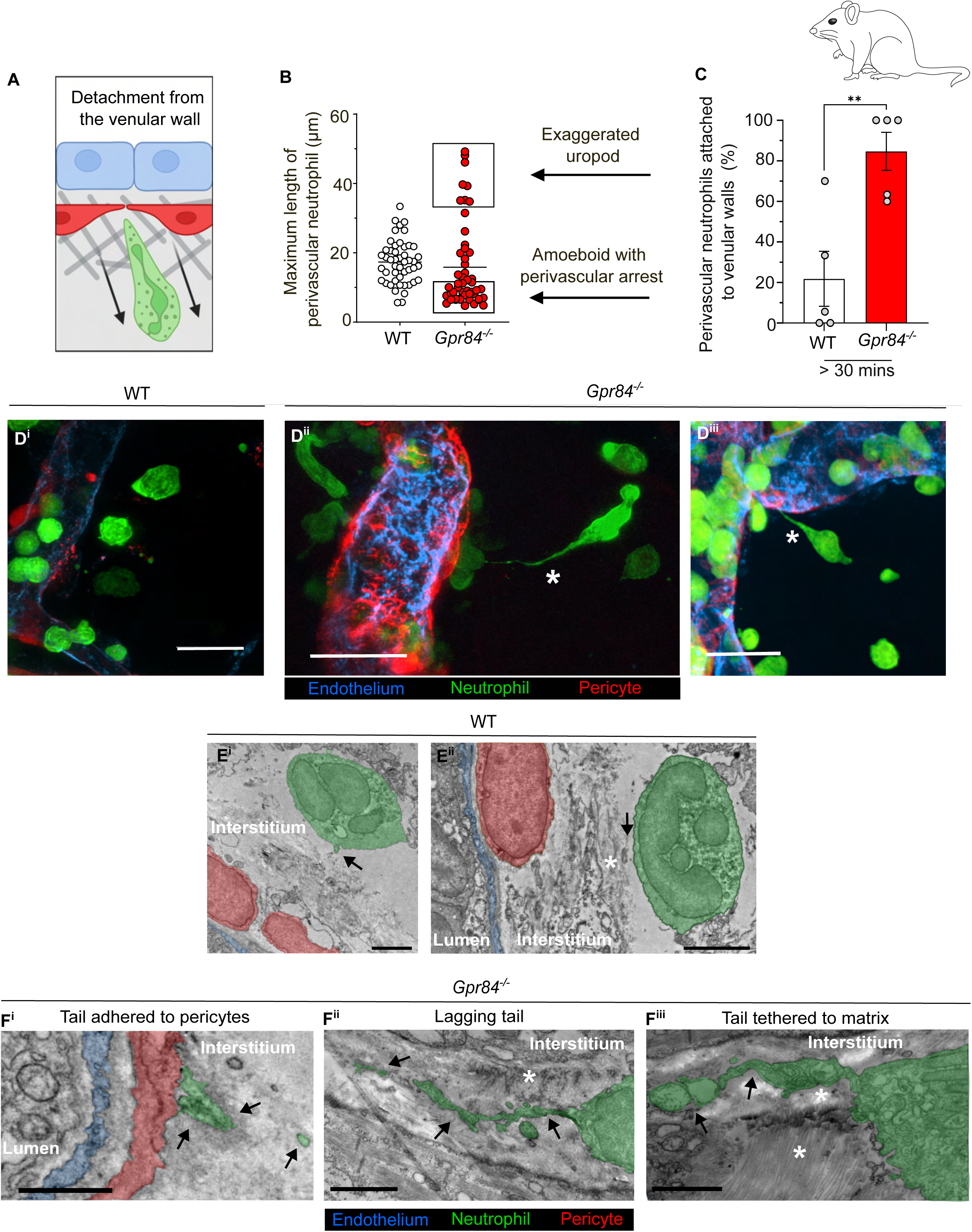
Neutrophil detachment from inflamed venular walls is impaired in GPR84-deficient mice. **(A)** Schematic depicting the response being analysed in Panels B-C, i.e. neutrophil detachment from the venular wall after breaching the pericyte layer. **(B)** The maximum length of an extravasated neutrophil whilst still tethered to the vascular wall (defined as perivascular neutrophil) during a 1 h recording of a TNF-stimulated post-capillary venule. Each data point represents an individual neutrophil pooled from WT and *Gpr84^-/-^* mice (n= 5 mice/ group). For the *Gpr84^-/-^* neutrophils, arrows indicate either an “exaggerated uropod” with an undulating motion prior to detachment or an “amoeboid” morphology with perivascular arrest. **(C)** Quantification of the percentage of perivascular neutrophils attached to the venular wall for > 30 min during the 1 h imaging period. **(D)** Representative images of neutrophils extravasating in WT (D’) and *Gpr84^-/-^* (D’’, D’’’) cremaster muscles. Cells were identified as follows: Neutrophils (*Lyz2-EGFP*; green), endothelial cells (anti-PECAM-1 mAb; blue) and pericytes (*Acta2-RFPcherry*; red). Asterisks indicate *Gpr84^-/-^* mice displaying a protracted attachment to the venular wall at the 4 h point of the reaction. **(E - F)** Transmission electron microscopy analysis of inflamed cremaster muscles showing the ultrastructure of neutrophils as they detach from the venular wall post-extravasation in WT **(E)** and *Gpr84^-/-^* **(F)** mice. **(E^i^ and E^ii^)** Representative images of WT neutrophils just after detaching from the venular wall at the experimental endpoint of 4 h. Black arrows indicate the remaining neutrophil (green) uropod post-detachment from the pericyte layer (red) and endothelium (blue). White asterisk indicates the presence of extracellular matrix components in the interstitial space. **(F)** Representative images of protracted neutrophil attachments to venular walls in *Gpr84^-/-^* mice, **(F^i^)** displaying retained pericyte attachment, **(F^ii^**) with a lagging tail, as indicated by arrows, and **(F^iii^)** tail retention along extracellular matrix. Data in C represents Mean ± SEM and analysed by an unpaired T-test (n=5 mice/group). ** P <0.01. Scale bars: D = 20 µm; E^i^ = 2 µm; E^ii^ = 3 µm; F = 1.5 µm.

### Transcriptomic analysis of GPR84-deficient murine neutrophils indicates defective actin cytoskeletal regulation and cell adhesion

With our live imaging studies strongly suggesting a role for Tre1/GPR84 in leukocyte detachment during extravasation, we next wished to gain insight into the likely signalling machinery involved. For this, we undertook an unbiased transcriptomic analysis of circulating WT and GPR84 deficient neutrophils. Principal Component Analysis (PCA) of data acquired from sorted blood neutrophils (isolated from untreated mice) showed a separation between WT and KO neutrophils (**Figure 6A**). This revealed 713 differentially expressed genes (DEGs), with 367 significantly upregulated and 346 significantly downregulated in GPR84-deficient neutrophils relative to WT cells **(Figure 6B**). Gene set enrichment analysis indicated dysregulated cytoskeletal remodelling and cell adhesion properties in GPR84-deficient neutrophils **(Figure 6C and 6D^i^-D^ii^)**, consistent with the observed phenotype of impaired detachment from the venular wall. Specifically, numerous candidates related to cytoskeletal remodelling were down-regulated in GPR84-deficient neutrophils **(Figure 6D^i^)**, including *Pak1*, *Arhgap5, Ccdc125* - all considered essential effectors of Rho-GTPases (Shin et al., 2013; Zhao and Manser, 2005). Interestingly, GPR84-deficient neutrophils also upregulated Rho regulators such as *Pip5k1*, a kinase implicated in RhoA polarisation and uropod formation in neutrophils (Xu et al., 2010).

**Figure 6.**
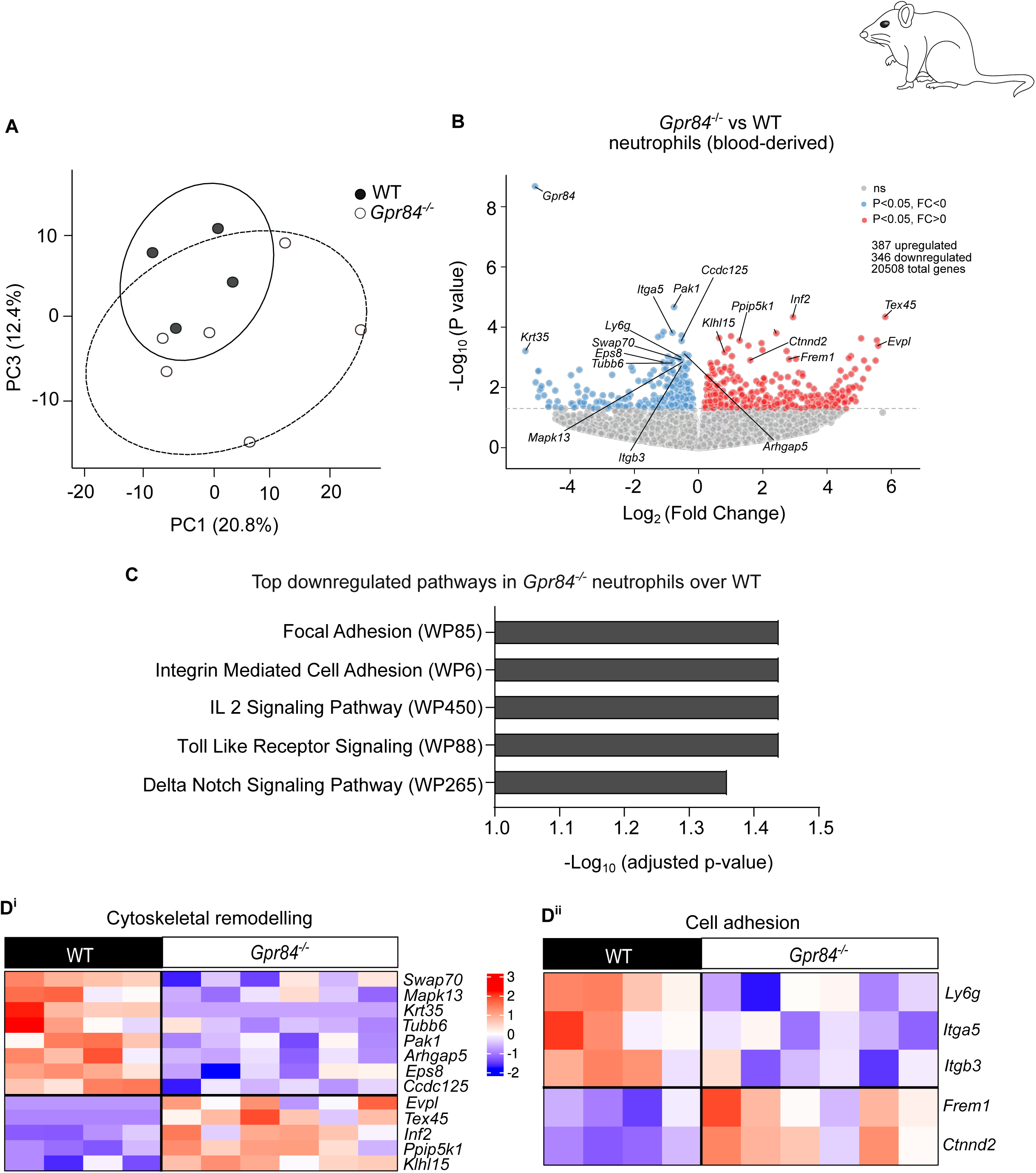
GPR84-deficient neutrophils express molecular changes in actin cytoskeletal regulation and cell adhesion. (A-D) Neutrophils were FACs sorted from the whole blood of WT (n=4) or *Gpr84^-/-^* (n=6) mice under baseline conditions and subjected to bulk RNA-sequencing. **(A)** Principal component analysis (PCA) of transcriptomes: PC1 and PC3 describe the major variance between WT (n=4) and *Gpr84^-/-^* (n=6) neutrophil transcriptomes. **(B)** Volcano plot showing the differentially expressed genes (DEGs) between WT and *Gpr84^-/-^* neutrophils with an unadjusted p-value of < 0.05. Labelled genes are associated with cytoskeletal remodelling and cell adhesion biological functions (also presented in heatmaps **(D)**. Significantly differentially expressed genes (unadjusted p-value> 0.05) are color coded as follows: genes downregulated in *Gpr84^-/-^* over WT neutrophils are indicated in blue (fold change, FC < 0) and upregulated genes in *Gpr84^-/-^* over WT neutrophils are indicated in red (FC > 0). **(C)** Pathway enrichment analysis (using the WikiPathways 2024 database from EnrichR) of 346 DEGs downregulated in *Gpr84^-/-^* over WT neutrophils (scored as −log_10_ (*p*-value)). The presented results are based on Fisher’s exact test with false discovery rate (FDR) adjustment. **(D)** Mini-heatmaps show transcript levels (with an unadjusted p-value <0.05) of genes associated with **(D^i^)** cytoskeletal remodelling and **(D^ii^)** cell adhesion, with the scale bar representing Z scaled gene expression across the samples.

Several DEGs indicative of altered cell adhesion, most notably integrins involved in cell-cell attachment and cell-ECM adhesion, were also identified **(Figure 6D^ii^)**. Specific examples that were down-regulated in KO neutrophils include *Itga5* that encodes for the α subunit of the integrin fibronectin receptor α5β1 (VLA-5) and *Itgb3* that encodes for the β3 integrin subunit. The latter is a component of multiple cell surface receptors that play roles in cell adhesion, migration and signalling and participates in RGD-dependent binding to ECM ligands (Chastney et al., 2025). Whilst suggesting defective adhesion of KO neutrophils to venular cells and basement membrane structures, we found no significant difference in the cell surface protein expression of key integrin subunits in KO vs WT neutrophils (**Supplementary Figure 3A**). The latter may be due to potential regulation of transcripts by negative feedback to maintain appropriate protein expression levels. KO neutrophils also showed altered expression of other genes implicated in cell migration such as *Ly6g,* a down-regulated candidate implicated in neutrophil infiltration and considered to act as a modulator of integrin functions (Song et al., 2025; Wang et al., 2012). Interestingly, KO cells displayed up-regulation of *Frem1* that encodes for generation of extracellular matrix proteins involved in formation of basement membranes (Smyth et al., 2004). Though not previously linked to neutrophils, its upregulation may be a cellular compensatory mechanism aimed at normalising impaired integrin-ECM interactions. Collectively, these data suggest that Tre1/GPR84 might regulate leukocyte extravasation via Rho-dependent cytoskeletal rearrangement and regulation of integrin adhesion dynamics.

### *Drosophila* Tre1 supports localized Rho1 activity during hemocyte extravasation

Based on our live *in vivo* imaging and transcriptomic data, we speculated that Tre1/GPR84 may regulate detachment of hemocytes/neutrophils from the vessel wall by Rho-mediated events. Tre1 contains two highly conserved sequences (the E/N/DRY and NPxxY motifs), known to act via distinct signaling pathways (G-protein signaling and Rho1/E-cadherin, respectively) for different elements of the spatio-temporal guidance of germ cells (LeBlanc and Lehmann, 2017). A sequence comparison of these motifs between *Drosophila*, mouse and human highlighted that the NPxxY (so called Rho binding domain), is fully conserved between Tre1 and mammalian GPR84, whereas the E/N/DRY is not (**Figure 7A**). *Drosophila* studies of migrating germ cells have shown the NPxxY motif is required for downstream Rho1 activation and that Tre1^NPxxY^ mutant germ cells demonstrate aberrant migration (Kim et al., 2021; LeBlanc and Lehmann, 2017). Our *in silico* AlphaFold3-mediated analysis (Abramson et al., 2024) of the predicted interface between *Drosophila* Rho1 and Tre1 suggested that Rho guanine nucleotide exchange factor 2 (RhoGEF2) and G protein alpha subunit o (Gαo) might act as intermediate accessory proteins to the stable structure (**Figure 7B**). RhoGEF2 is an established activator of Rho1 that acts by facilitating the switch of GDP to GTP (Mosaddeghzadeh and Ahmadian, 2021). Since we found hemocytes and neutrophils lacking Tre1/GPR84 to exhibit similar morphological phenotypes of lagging tails and transcriptomic alteration of Rho family members, we hypothesized that these GPCRs may signal through their conserved NPxxY motifs. Consequently, we considered that the latter would then activate Rho1 via RhoGEF2 and Gαo to facilitate tail retraction/detachment during the latter stages of extravasation (**Figure 7C**).

**Figure 7:**
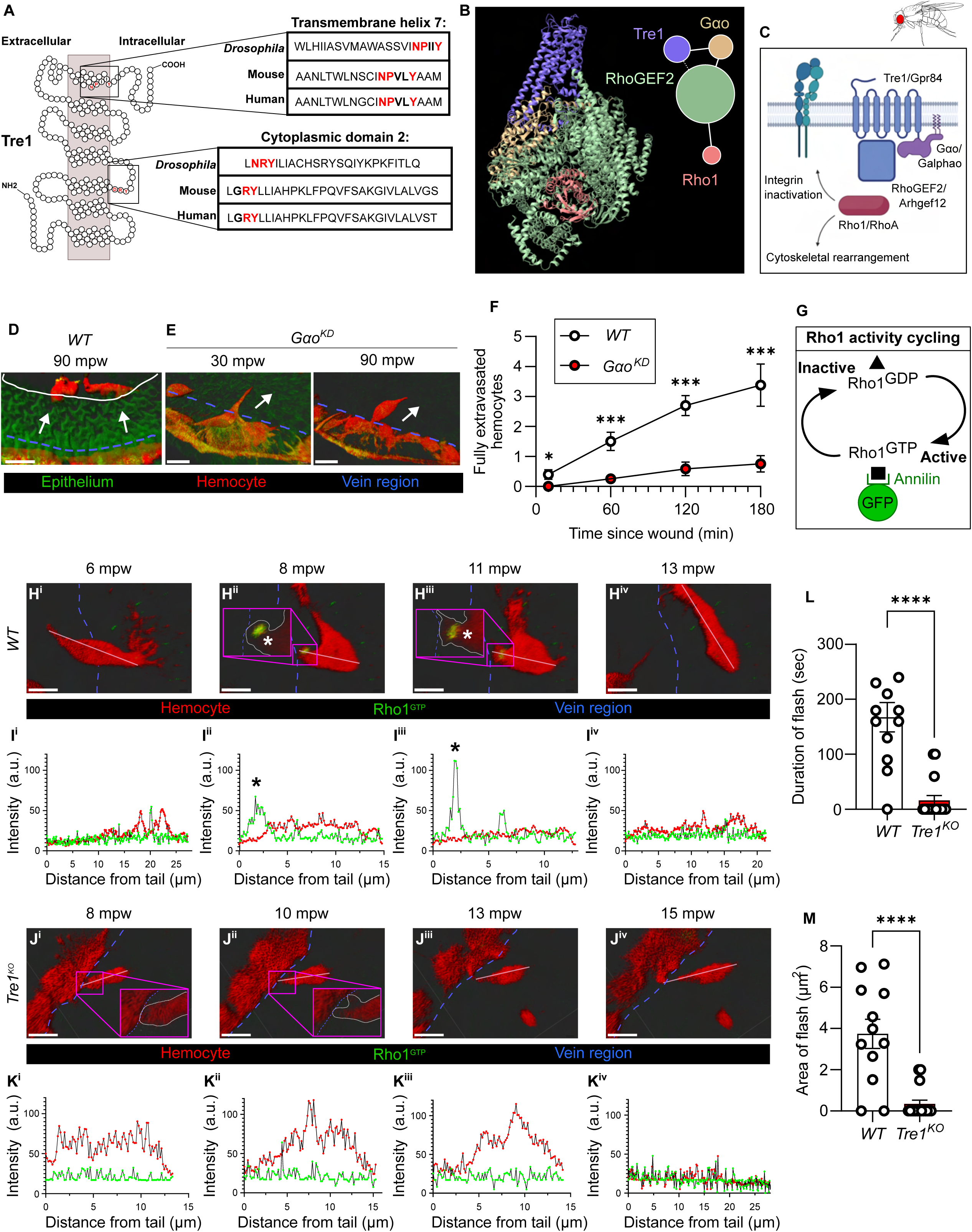
During extravasation, Tre1 signaling supports dynamic localization of active Rho1 to hemocyte lagging tails during tail retraction. **(A)** Pearl necklace diagram of Tre1 highlighting key motifs, and how these are conserved between *Drosophila*, mice and humans. **(B)** Structural model generated in Alphafold3 showing the predicted interface between Tre1 (blue) and Rho1 (red), with RhoGEF2 (green) and Gαo (brown) acting as intermediate accessory proteins). Visualization rendered in ChimeraX. **(C)** Proposed signaling pathway linking Tre1 to hemocyte tail retraction from the vasculature. Created in BioRender. L, C. (2025) https://BioRender.com/bup6fba. **(D - E)** IMARIS renderings of time-lapse confocal imaging demonstrating the requirement of Gαo for hemocyte (red) extravasation from the vasculature (blue dotted line) towards wounds (solid white outline). (D) *WT* hemocytes are recruited towards wounds. (E) *Gαo^KD^* hemocytes phenocopy Tre1 mutants with apparent vessel de-adhesion defects. **(F)** Quantification of numbers of extravasation events in *WT* versus *Gαo^KD^* hemocytes. **(G)** Diagram showing how the Annilin active-Rho1 biosensor reports Rho1 activity. **(H - J)** IMARIS renderings and intensity line plots of *in vivo* time-lapse imaging of *WT* (H^i^ - H^iv^) and *Tre1^KO^* (J^i^ - J^iv^) hemocytes (red) as they extravasate from vessels, with active Rho1 (green). **(I^i^ - I^iv^)** In *WT* hemocytes a transient puncta of active-Rho1 localizes in the tail that resolves during tail retraction; black and white asterisks. **(K^i^ - K^iv^)** In *Tre1^KO^* hemocytes there are no active Rho1 flashes. **(L)** Quantification of the time taken for active Rho1 flashes to resolve and **(M)** area of flashes between genotypes. Data in line plot **(F)** were analyzed with multiple unpaired T-tests followed by correction for multiple comparisons using the two-stage step-up method of Benjamini, Krieger, and Yekutieli (FDR control). Data in bar plots **(L and M)** represent calculated mean for each genotype, and dots represent individual hemocytes pooled from multiple animals. For each quantification data represents at least 7 animals per any genotype and represented is the mean ± SEM. Asterisks = *P<0.05, **P<0.01, ***P<0.001, ****P<0.0001. Scale bars: D, E, H^i^ - H^iv^ and J^i^ - J^iv^ = 10 µm. See also **Supplementary Movie 7**.

To address this proposed mechanism, we utilised high-resolution *in vivo* imaging of extravasating *Drosophila* hemocytes in response to a laser wound. Analysis of Gαo-RNAi knockdown hemocytes revealed Gαo to be essential for timely and proper hemocyte extravasation from the vasculature. Indeed, Gαo-RNAi knockdown phenocopied Tre1 mutants with hemocytes displaying de-adhesion defects (**Figure 7D - F, Supplementary Figure 4A - B** and **Supplementary Movie S6**). Utilizing a transgenic fluorescent biosensor for *in vivo* Rho1 activity that consists of GFP tagged to the Rho1 GTP-binding domain of Anillin (Munjal et al., 2015) we visualized the dynamics of Rho1 activity *in vivo* during extravasation (**Figure 7G - M** and **Supplementary Movie S7**). As control hemocytes extravasated from vessels, we observed clear punctae of active Rho1 localized to the hemocyte tail (preceding tail retraction) that dissolved after the tail had detached (**Figure 7H - I**). In Tre1-deficient hemocytes, active Rho1 was absent from the tail, which remained adherent for longer and often failed to fully detach from the vein (**Figure 7J - K**). Quantification of Rho1 activity (duration and area of “active Rho1” fluorescence) confirmed that Tre1-null hemocytes have smaller and more transient flashes of localized Rho1 activity, even on the rare occasions when they occurred (**Figure 7L - M**).

### Leukocyte Tre-1/GPR84 modulate integrin-mediated adhesion

A key finding of the present study is that hemocytes and neutrophils lacking Tre1/GPR84 have long lagging tails that can persist for hours after they have attempted to leave the circulation. In control mammalian leukocytes, uropod adhesion and then retraction results from the accumulation of integrin-to-ECM focal adhesions that dissolve through a combination of contractile and biochemical forces (Hind et al., 2016). We speculated that Tre1/GPR84 facilitate tail retraction through regulation of cytoskeletal contraction and dissolution of integrin focal adhesions mediated by Rho1/A activation. This notion is supported by previous *in vitro* studies showing that pharmacological blockade of RhoA leads to abnormally elongated tails in monocytes adherent to extracellular matrix (Worthylake et al., 2001), an observation that resembles Tre1/GPR84 null immune cells attempting to extravasate *in vivo*.

In hypothesizing that activated Tre1 dissolves integrin adhesions in the hemocyte tail, we predicted that elevating hemocyte integrin levels would phenocopy Tre1-deficient hemocytes. Indeed, we observed that overexpression of *myospheroid* (*mys*; the predominant *Drosophila* β-integrin) specifically in hemocytes led to a strikingly similar defective extravasation phenotype (**Figure 8A - D**). To further address whether Tre1/GPR84 regulates focal adhesion dissolution in lagging tails as leukocytes extravasate, we examined whether *mys* levels were perturbed in *Drosophila* lacking Tre1. We found that in Tre1 null pupae, *mys* mRNA levels were significantly reduced, as were transcripts for *focal adhesion kinase* (*Fak*) a known regulator of integrins (**Figure 8E - F**). These data are consistent with the transcriptomic analysis of GPR84 KO mouse neutrophils, suggesting a conserved role for Tre1/GPR84 in modulating integrin-based adhesion **(Figure 6C),** and our observations of defective adhesion of GPR84 mouse KO neutrophils to intercellular adhesion molecule 1 (ICAM-1)-coated plates (**Figure 8G - I**). ICAM-1 is highly expressed on TNF-stimulated endothelial cells and pericytes (Proebstl et al., 2012; Cooper et al., 2024; Recio et al., 2018; Yousefi et al., 2001; Bouchard et al., 2007) and was thus used as an *in vitro* adhesive substrate to functionally mimic neutrophil interactions with vascular cells. Following stimulation with the chemokine CXCL1, *Gpr84^-/-^* neutrophils displayed a significantly reduced **“**spreading” morphology and an overall defective motility *in vitro* (**Figure 8G - I**). These defects were associated with reduced membrane ruffling and polarization, compared to controls, suggesting an impairment in integrin-induced cytoskeletal remodeling in response to activation (**Figure 8G - I**). Collectively, the data acquired from our *Drosophila* and mouse studies, using both *in vitro* and *in vivo* assays, strongly support the concept that Tre1/GPR84 exhibit conserved functionalities in supporting Rho-mediated regulation of integrin activity.

**Figure 8:**
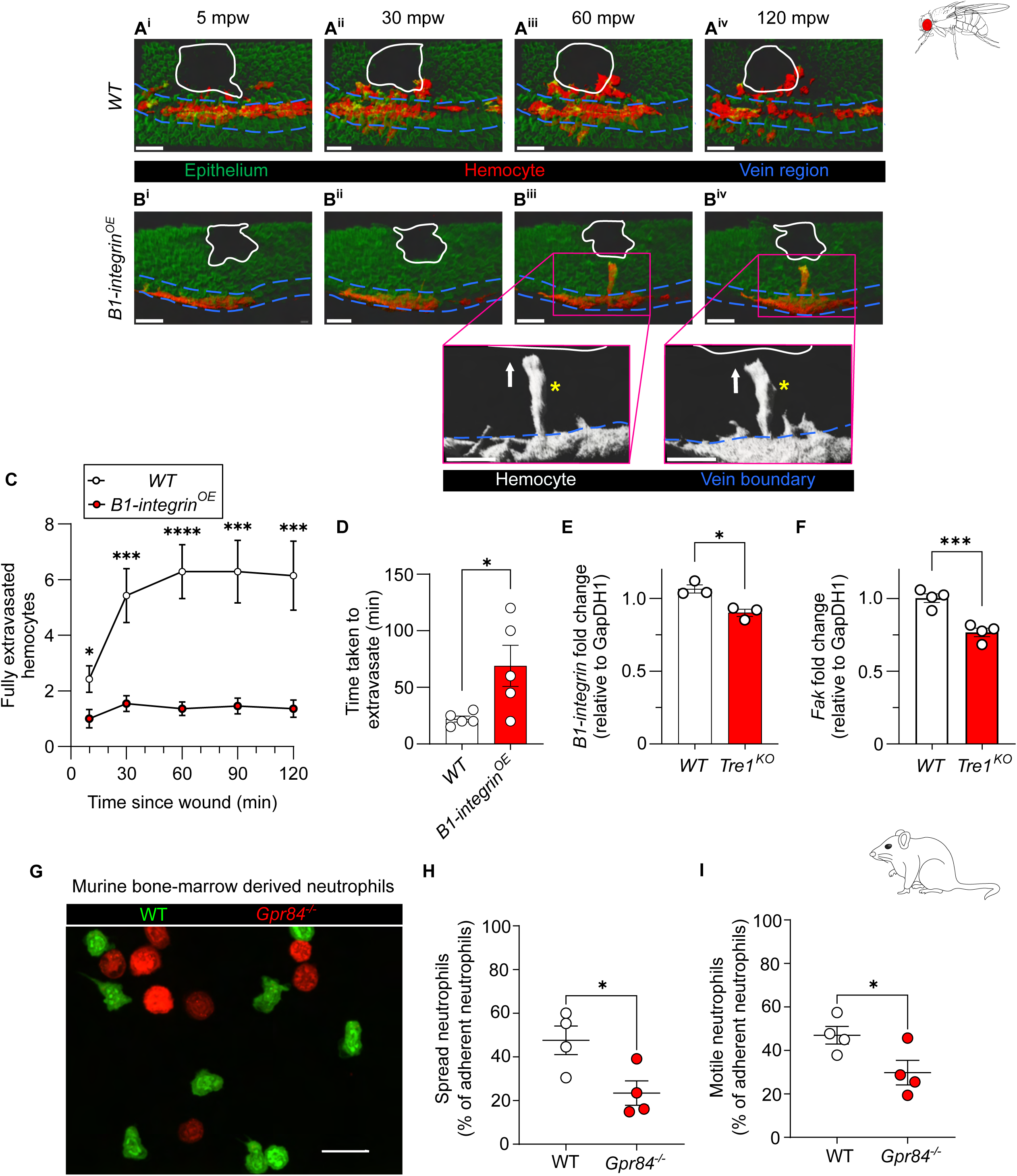
Tre1/GPR84 regulate integrin expression and functions. (A and. **B)** IMARIS rendering of *in vivo* imaging of WT versus *myospheroid* (B1-integrin) over-expressing hemocytes as they respond to wounds, with labelling of epithelium (green, *ubi-GMA*), hemocytes (red, *srp-Gal4>mCherry, srp-mch*), veins (manually labelled with a blue dotted line) and wound perimeter (solid white line). **(A^i^ - A^iv^)** *WT* hemocytes are recruited to the wound and fully detach from the vasculature **(B^i^ - B^iv^),** whereas B1-integrin**^OE^** hemocytes show reduced recruitment, a lagging tail phenotype with long protrusions (yellow asterisks) towards the wound, tethering to the vasculature. **(C)** Quantification of numbers of hemocytes that have fully dissociated from veins and extravasated towards wounds. **(D)** Analysis of time taken for individual hemocytes to extravasate from veins in both *WT* and B1-integrin**^OE^** genotypes. **(E and F)** Quantitative RT-PCR of the relative expression of *B1-integrin* and *Fak* in whole 75h APF pupae in both *WT* and *Tre1^KO^* backgrounds, with both target genes’ threshold cycle values normalized to Gapdh1; in the *Tre1^KO^* background both genes show a significant decrease in expression levels. RT-PCR data represented in bar plot is Mean ± SEM overlayed with average value (red dots) for each biological replicate (RNA pooled from 10 individuals). **(G-I)** Bone marrow-derived WT (green) and *Gpr84^-/-^* (red) mouse neutrophils were mixed and seeded onto ICAM-1-coated surfaces and stimulated with CXCL1 for 15 mins. **(G)** Representative confocal images illustrating the cellular morphology of WT and KO neutrophils, and quantified proportion of **(H) “**spread” and **(I)** motile adherent neutrophils. Data shows Mean ± SEM. **(D, E, F, H and I)** Data was analysed by unpaired T-tests. Scale bar: **A^i^ - A^iv^** and **B^i^ - B^iv^** = 15 µm; **G** = 20 µm.

### Pharmacological blockade of GPR84 suppresses neutrophil extravasation in acute inflammation

Given the conserved role for GPR84/Tre1 in supporting effective immune cell extravasation, in a final series of experiments we explored the effects of pharmacologically modulating murine GPR84 during acute inflammation. For this purpose, we tested the efficacy of a selective and orally active functional GPR84 antagonist (GLPG1205) in multiple murine models of acute inflammation. Initially, using our TNF-inflamed cremaster muscle model, a reaction in which the GPR84 KO mice exhibited ∼70% inhibition of leukocyte migration (**Figure 3D**), we found that treatment of mice with the antagonist led to a dose-dependent suppression of neutrophil extravasation **(Figure 9A - B)**. Using the higher dose of the antagonist (30mg/kg), these findings were extended to three other inflammatory settings, namely a zymosan-induced peritonitis model, a cutaneous ischemia-reperfusion (IR) injury model and finally a skin wounding assay. In the peritonitis model, a well-characterised self-resolving acute inflammatory reaction, 24 hrs post injection of zymosan, GLPG1205 pre-treated mice exhibited reduced peritoneal cavity infiltration of both neutrophils and monocytes (**Figure 9C**). These results suggest that GPR84 can mediate appropriate migration of multiple populations of myeloid cells. The cutaneous IR injury model, employed as a model of a pressure ulcer, similarly showed an effective neutrophil infiltration response in control mice that was significantly suppressed in mice treated with the GPR84 antagonist (**Figure 9D**). The latter studies included both male and female mice, demonstrating that the efficacy of GPR84 blockade was not sex specific.

**Figure 9.**
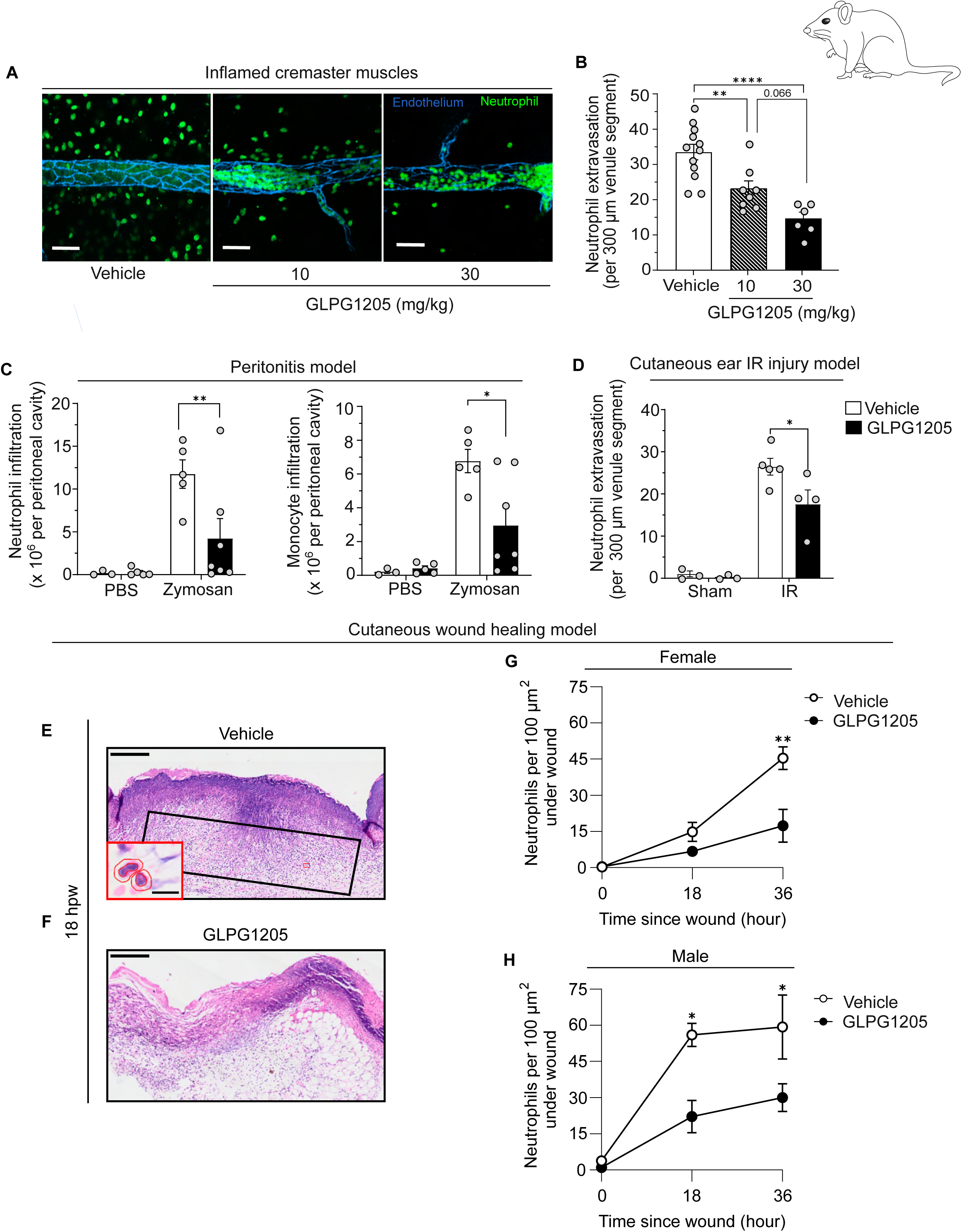
Pharmacological blockade of GPR84 suppresses neutrophil extravasation in models of acute inflammation. **(A)** Representative images of TNF-stimulated (4 h) cremasteric venules of C57Bl6/J mice pre-treated (oral gavage, 24 h before) with vehicle or GLPG1205 (10 or 30 mg/ kg). Tissues were fixed, permeabilised and stained for endothelial cells (PECAM-1; blue) and neutrophils (MRP-14; green). **(B)** Quantification of the average number of extravasated neutrophils per 300 µm venule segment from 5-10 venules per mouse; n= 6-12 mice/group. **(C)** C57BL/6 mice were treated with vehicle or GLPG1205 (30 mg/ kg) 24 h prior to an i.p. injection of zymosan or PBS control. The infiltration of neutrophils and monocytes at 24 h post injection was quantified by flow cytometry (n = 5 – 8 mice/group). **(D)** C57BL/6 mice were treated with vehicle or GLPG1205 (30 mg/ kg) 24 h prior to ear ischaemia reperfusion injury. Quantification of the average number of extravasated neutrophils from 6-10 venule segments per mouse (with n = 3-5 mice / group). **(E and F)** Haematoxylin and Eosin stained wax sections of back skin wounds at 18 h post wounding (hours post wounding, hpw). **(E)** Representative image of an 18 h wound from a vehicle treated female mouse. Black box indicates approximate region for neutrophil quantification and red box indicates site of inset highlighting automated capture and quantification of numbers of neutrophil nuclei. **(F)** Representative image of a 18 h wound from a GLPG1205 treated female mouse. **(G)** Quantification of neutrophil extravasation at different time-points in the wound beds of vehicle versus GLPG1205 treated female **(G)** and male **(H)** mice. Data shows Mean ± SEM. Data in **(B)** was analysed by one-way ANOVA with a Tukey’s test for multiple comparisons and **(C and D)** were analysed by Two-way ANOVA with Fisher’s LSD or Šídák’s multiple comparisons post hoc test. **(G and H)** were analysed by unpaired T-tests followed by correction for multiple comparisons using Šídák’s post hoc test. * P < 0.05, ** P <0.01. Scale bars: A = 50 µm; E and F = 250 µm: E inset 10 µm.

GPR84 expression increases in murine myeloid cells (neutrophils, macrophages and monocytes) following skin wounding (Cooper et al., 2024). To investigate how wound inflammation might be impacted by GPR84 signalling, we generated punch biopsy skin wounds in GLPG1205-treated versus control adult male and female mice (**Figure 9E - H**). Standardly, the initial influx of neutrophils to such a skin wound begins from 6 h post wounding, with numbers peaking at 24 h and diminishing back to basal levels after 48 h, as macrophage numbers begin to predominate in the wound. To capture potential differences in neutrophil influx in treated versus control mice, we harvested wounds at 18 and 36 h and analysed wound sections stained with Hematoxylin and Eosin (**Figure 9E - F**); here, the automated software QuPath (Bankhead et al., 2017) enabled efficient quantification of Hematoxylin-stained neutrophil nuclei within Eosin-stained cytoplasms (inset, **Figure 9E**). Our data show a clear dampening of neutrophil influx into the wound bed at both timepoints in female (**Figure 9G**) and male (**Figure 9H**) mice.

Collectively, using a pharmacological approach, these results demonstrate the importance of GPR84 in regulating leukocyte extravasation in several inflammatory contexts in different organs, in response to different stimuli and in both male and female animals.

## Discussion

Whilst the molecular and cellular pathways that regulate the onset of leukocyte extravasation across vessel walls are increasingly well characterized, the complex signaling events that enable successful completion of the sequential steps in vessel breaching remain unclear. Here, we identified the orphan GPCR, Tre1/GPR84, as pivotal for effective termination of leukocyte extravasation during acute physiological and pathological inflammation. Specifically, by revealing how leukocytes deficient in these GPCRs retain long, adherent tethers and fail to complete extravasation *in vivo,* we demonstrate their functional role in effective leukocyte tail detachment from vessel walls. Crucially, our murine studies demonstrated a role for GPR84 in multiple stages of neutrophil extravasation, consistently indicative of regulating neutrophil de-adhesion from vascular cells, endothelial cells and pericytes, as well as potentially the venular basement membrane. Mechanistically, transcriptomic analyses together with intravital imaging of subcellular signaling events revealed that Tre1/GPR84 regulated leukocyte cytoskeletal and adhesion dynamics, specifically supporting localized Rho1 activity in retracting leukocyte tails. Moreover, pharmacological antagonism of murine GPR84 *in vivo* dampened leukocyte infiltration in a range of acute inflammatory conditions in both sexes, including zymosan-induced peritonitis and skin wounding. Collectively these data identify GPR84 as a previously unknown regulator of neutrophil extravasation and present data to suggest its potential as a promising anti-inflammatory therapeutic target.

GPR84 is widely considered as a pro-inflammatory molecule as its expression is increased in myeloid cells (particularly monocytes and macrophages) in response to multiple inflammatory stimuli (Cooper et al., 2024; Recio et al., 2018; Yousefi et al., 2001; Bouchard et al., 2007; Nagasaki et al., 2012). Furthermore, studies have reported on the ability of GPR84 to mediate leukocyte infiltration into inflammatory sites and indeed this receptor has been the subject of much interest as a therapeutic anti-inflammatory target (Cooper et al., 2024; Recio et al., 2018; Suzuki et al., 2013; Wang et al., 2023b; Mancini et al., 2019). Despite the significant attention given to this molecule for >15 years, the mechanism through which it mediates leukocyte trafficking has remained unknown. Here we demonstrate a specific role for Tre1/GPR84 in leukocyte (*Drosophila* hemocyte and murine neutrophil) extravasation across vessel walls in acute inflammation. We propose that Tre1/GPR84 signaling acts via the G protein alpha subunit O (GaO) to promote dynamic Rho1 activation within the leukocyte tail during extravasation; Rho1 activity in turn promotes cytoskeletal and adhesion remodeling that enable the leukocyte to fully detach its lagging tail from the vessel wall. In the absence of effective Tre1/GPR84 signaling, despite the leukocyte initiating the process of extravasation, the lack of Rho1-driven de-adhesion leads to decelerated breaching of vessel walls, culminating in a striking elongated leukocyte phenotype in the perivascular region. The resultant impeded extravasation led to a notable accumulation of ‘stuck’ leukocytes around the inflamed vessel. Interestingly, despite leukocytes exhibiting a more adhesive cellular phenotype, we found a marked transcriptomic downregulation of adhesion molecules (particularly integrin subunits) within Tre1/GPR84 deficient leukocytes. This raises the possibility that null leukocytes downregulate adhesion-related gene expressions to compensate for the elevated adhesion response.

Though it is widely accepted that tightly controlled spatio-temporal regulation of integrin-mediated leukocyte adhesion is essential for effective extravasation (Nourshargh and Alon, 2014), the precise associated mechanisms remain unclear. Integrins play established roles in driving the initial steps of neutrophil extravasation, with activation of defined β2 integrins mediating distinct intraluminal responses, including LFA-1-dependent adhesion followed by Mac-1-dependent crawling (Phillipson et al., 2006). Additionally, there is ample evidence for the involvement of integrins in supporting neutrophil breaching of vascular components, ECs, pericytes and the venular basement membrane (Nourshargh and Alon, 2014; Girbl et al., 2018). Finally, prior to full exit from the venular wall, neutrophils (and other immune cells) exhibit elongated uropods in the perivascular space, a phenomenon mediated through integrin-mediated attachment of immune cells to vascular structures (Hyun et al., 2012); here, it has been shown that a coordinated dissociation of LFA-1 from the endothelium and VLA-3-mediated migration through the basement membrane enables timely de-adhesion and propulsion into the interstitial tissue. Our findings enhance our knowledge of these processes by presenting evidence for Tre1/GPR84 as an essential component of leukocyte de-adhesion from vessel walls, potentially inducing the dissolution of β2 integrin complexes via Rho-signaling at the terminal stages of neutrophil extravasation.

Despite improved understanding of the intracellular events downstream of activated Tre1/GPR84, the identity of the upstream signals mediating leukocyte GPR84 activation (including its precise ligand) remain unclear. Studies of the *Drosophila* germ cells reveal that the Wunens, germ cell-expressed membrane proteins that can dephosphorylate and inactivate phospholipids, may modulate Tre-1 signaling by creating a phospholipid gradient (Chen et al., 2025). In the mammalian system, GPR84 has been proposed as a medium-chain fatty acid (MCFA) receptor, whereby MCFAs promote leukocyte migration, phagocytosis and release of pro-inflammatory cytokines (e.g. TNF) (Suzuki et al., 2013; Nagasaki et al., 2012; Wang et al., 2006; Hu et al., 2008). Intriguingly, our transcriptomics data revealed an upregulation of the dual functional inositol kinase Ppip5k1 in *Gpr84* deficient neutrophils. *Drosophila* dPip5k generates PI(4,5)P2 and plays important role in polarizing the actin cytoskeleton for directed migration of embryonic germ cells, in cooperation with Tre1 (Kim et al., 2021). Signaling via phospholipids may thus play conserved roles modulating leukocyte Tre1/GPR84 activity in acute inflammatory settings *in vivo*.

Comparative transcriptomic profiling of control versus GPR84-deficient neutrophils revealed numerous biological pathways that could be regulated by GPR84. Whilst cytoskeletal remodeling and cellular (focal and integrin-mediated) adhesion pathways were amongst the most downregulated pathways in GPR84 KO cells – consistent with our data suggesting leukocyte Tre1/GPR84 supports Rho1-dependent tail detachment - neutrophils lacking GPR84 also downregulated other key immune pathways (including IL2 and Toll Like Receptor Signaling). IL2 signaling within neutrophils has been suggested to inhibit their migration, increase their adherence to ECs and stimulate neutrophil antimicrobial activities, including the respiratory burst and degranulation (Li et al., 1996; Kowanko and Ferrante, 1987). Our transcriptomic data show that components of the Delta/Notch signaling pathway were also downregulated by GPR84 deficiency. Whilst the role of Notch signaling in myeloid lineages (particularly neutrophils) remains poorly understood, evidence suggests Toll Like Receptor (TLR) signaling can induce Notch receptor and ligand expression and these pathways may coordinate to fine-tune leukocyte activation and function (Hildebrand et al., 2018; Hu et al., 2008). Notch signaling has also been linked to myeloid cell development and differentiation (Schroeder and Just, 2000), suggesting GPR84 may modulate leukocyte development, consistent with our observed downregulation of *Ly6g* (a candidate reflective of neutrophil maturity) in GPR84 deficient neutrophils. These data suggest GPR84 may have additional functions in leukocytes beyond Rho1-mediated regulation of leukocyte adhesion.

The human genome encodes more than 800 GPCRs that orchestrate diverse physiological processes, including key roles in leukocyte recruitment, activation and resolution (Sun and Ye, 2012). Leukocytes express multiple GPCR-type chemokine receptors (e.g. CCR5, CXCR2, CXCR4) that detect chemokine gradients produced by damaged, inflamed and infected tissues (Sun and Ye, 2012). GPCR activation triggers intracellular signaling that regulates leukocyte chemotaxis and key effector functions (such as degranulation and phagocytosis). Given that GPCRs are major therapeutic targets – accounting for nearly 35% of all clinically used drugs – GPCR antagonists are considered as plausible strategies for selective modulation of leukocyte trafficking in numerous inflammatory conditions. In line with this, there exists considerable therapeutic interest in GPR84 as a potential druggable target for various inflammatory conditions, despite its pathophysiological functions and molecular mechanisms remaining largely unknown. Here our data adds important mechanistic insight into GPR84’s translational potential. Our *in vivo* pharmacological studies revealed the efficacy of a GPR84 functional antagonist (GLPG1205) in dampening neutrophil infiltration in multiple experimental disease models, including skin wounding and peritonitis. This is consistent with recent work identifying GPR84 as a key regulator of myeloid cell (particularly macrophage) recruitment during murine skin wound healing (Cooper et al., 2024). Intriguingly, GPR84 levels are also increased in murine models of diabetes and fibrosis (Gagnon et al., 2018; Recio et al., 2018), with inhibition of GPR84 signaling reducing fibrosis in a range of tissues, including the skin, heart, lung, kidney and pancreas (Simard et al., 2020; Puengel et al., 2020; Gagnon et al., 2018). Moreover, initially identified within a myeloid cell signature specific to a zebrafish model of thermal injury, GPR84 has also been identified as a biomarker of burn injury exhibiting increased expression on blood neutrophils isolated from human patients (Hou et al., 2024). These studies, together with our work, indicate how GPR84 signaling might contribute to numerous inflammatory pathologies, including chronic non-healing wounds, peritonitis, acute respiratory distress syndrome, gastroesophageal reflux disease and ulcerative colitis (Abdel-Aziz et al., 2016; Yin et al., 2020; Zhang et al., 2022).

In summary, we have exploited the unique collective tractability of our *Drosophila* and murine models for rapid mechanistic interrogation of complex cell-specific signaling networks *in vivo.* With the focus of these studies being on regulation of leukocyte extravasation across vessel walls, integrating our *Drosophila* studies with murine models that enable intravital imaging of mammalian leukocyte extravasation has proven to be a powerful approach for identifying new, conserved and fundamental molecular players of acute inflammation. Capitalizing on *Drosophila’s* unrivalled potential for integrating *in vivo* real time imaging of subcellular signaling events with cell type-specific genetics, we have identified a conserved role for GPR84 orthologues in mediating localized leukocyte de-adhesion and cytoskeletal dynamics during extravasation. In the long-term, we envision the unique imaging and large-scale screening opportunities of *Drosophila* will enable the discovery of more genes with important conserved regulatory functions in immune cell extravasation, both within immune cells and the vascular cells with which they interact.

## Supplemental Figure Legends

**Figure S1.**
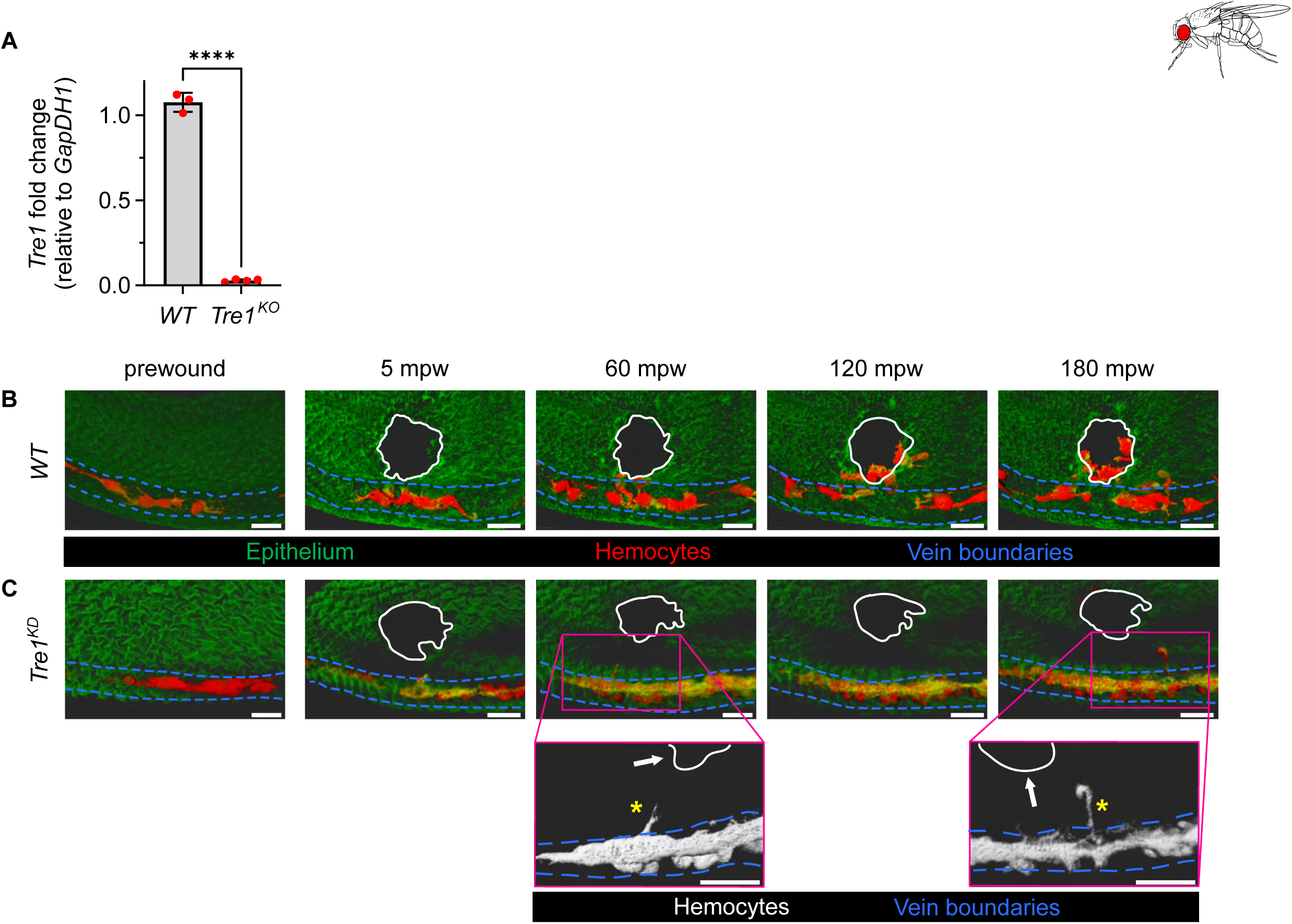
Tre1 is required for complete hemocyte extravasation. **(A)** Quantitative RT-PCR of the relative expression of *Tre1* in whole 75h APF pupae in *Oregon R* (*WT*) versus *Tre1^KO^* backgrounds, with *Tre1* threshold cycle values normalized to Gapdh1 validating the *Tre1^KO^* background. RT-PCR data represented in bar plot (whiskers, maximum and minimum values) overlayed with average value (red dots) for each biological replicate (RNA pooled from 10 individuals). **(B** and **C)** Representative *in vivo* time-lapse imaging of 75h APF of RNAi-mediated silencing of Tre1 within hemocytes (*srp-Gal4*-driven knockdown) results in a significant reduction in hemocyte extravasation capacity. Solid white line indicates border of wound, and arrows indicate direction towards wound that extravasated hemocytes travel. Scale bars: B and C = 15 µm.

**Figure S2.**
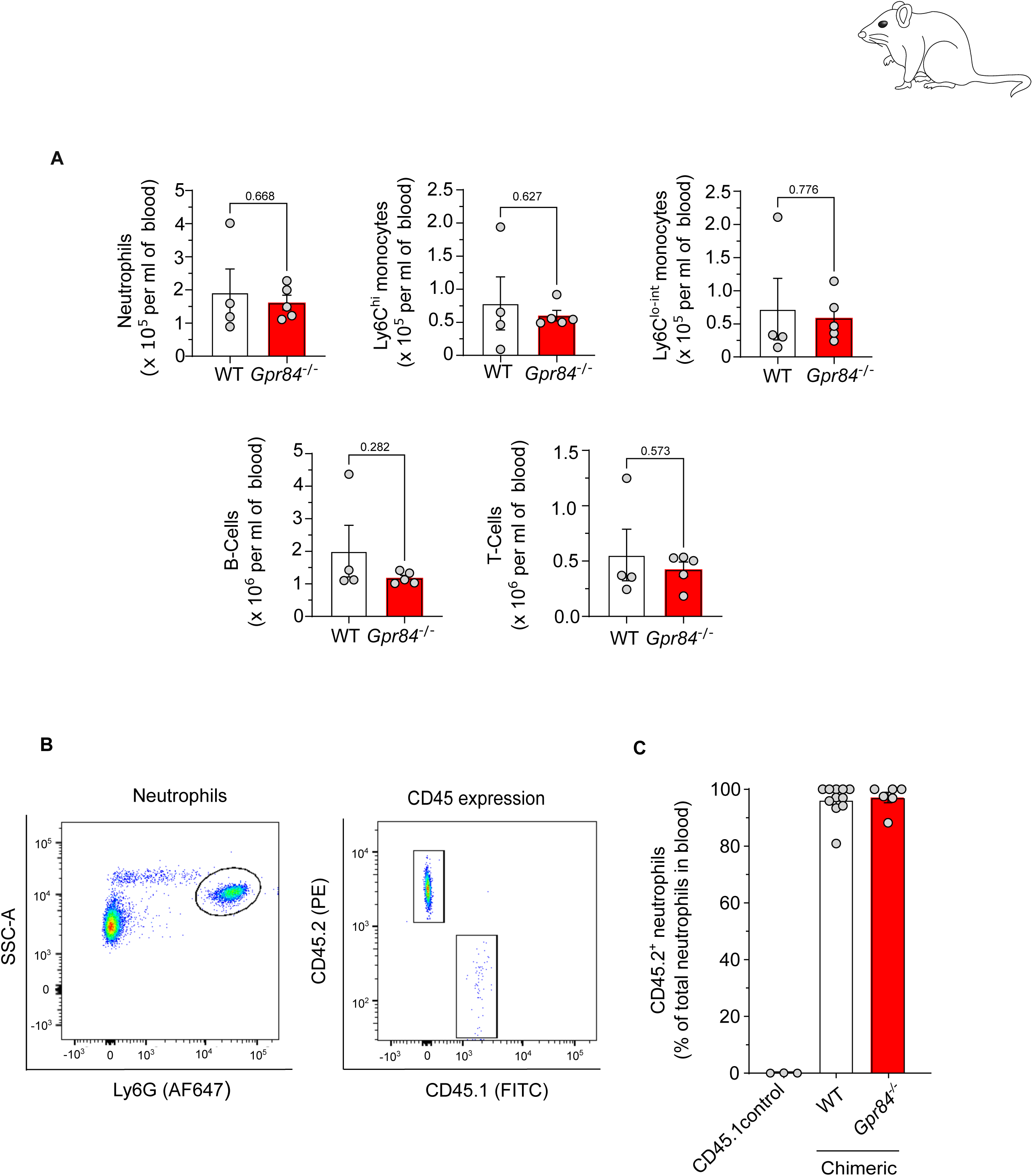
Blood leukocyte counts of WT vs *Gpr84^-/-^* global knockouts and reconstitution efficiency in chimeric mice after bone marrow transfer. (**A**) Differential leukocyte counts of whole blood from WT (n=4) and *Gpr84^-/-^*(n=6) mice determined by flow cytometry. **(B)** Representative flow cytometry gating strategy for selecting neutrophils, and **(C)** assessing CD45.2 allelic variant expression as a marker of reconstitution efficiency when generating WT and *Gpr84^-/-^* chimeric mice. Data represents Mean ± SEM and in **(A)** analysed by an unpaired T-test with no significance between experimental groups.

**Figure S3.**
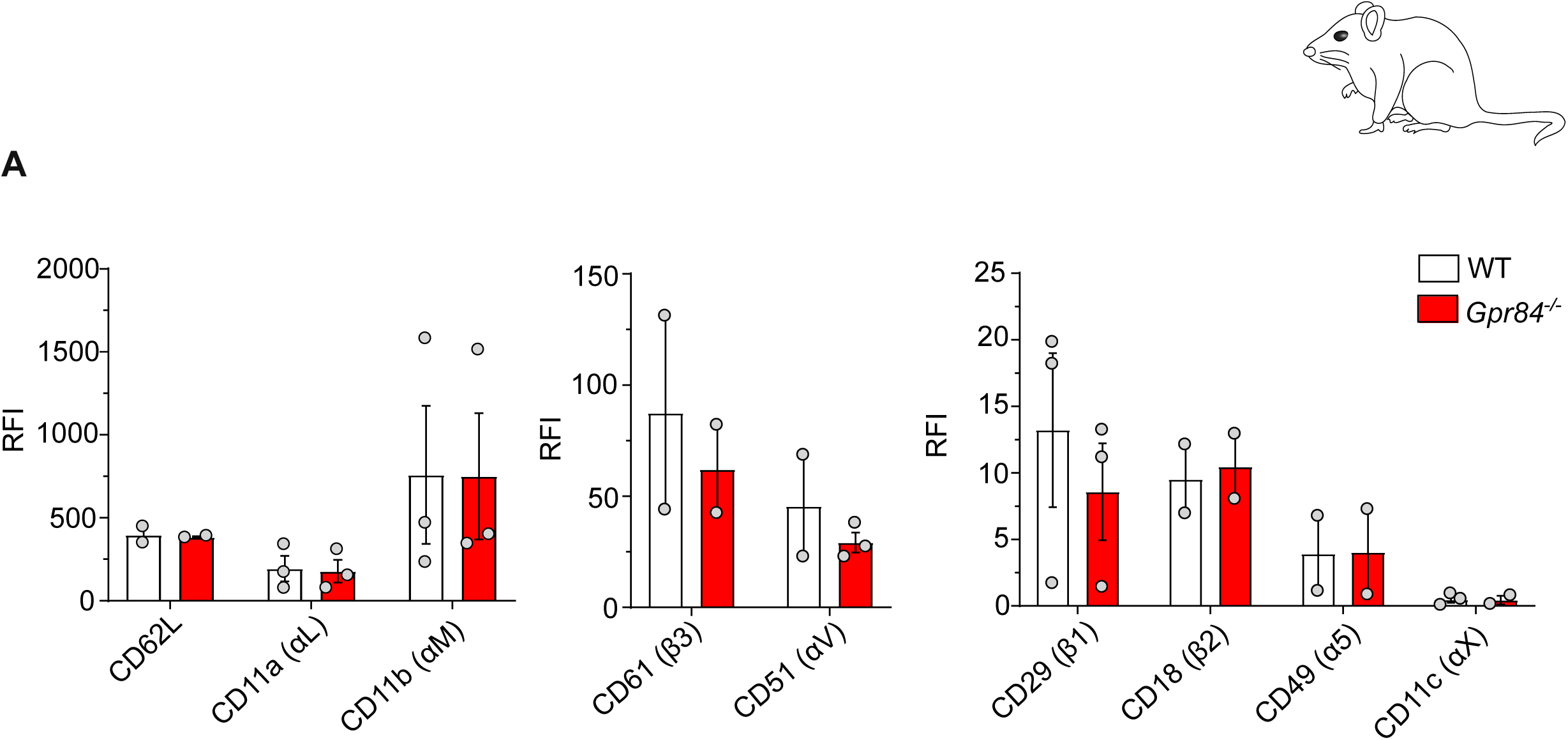
Integrin expression on WT and *Gpr84^-/-^* mouse blood neutrophils. **(A)** Relative fluorescence intensity (RFI) of surface expression levels of indicated integrins on mouse blood neutrophils, from WT and *Gpr84^-/-^* as determined by flow cytometry (2-3 mice/ group). Data represents Mean ± SEM.

**Figure S4.**
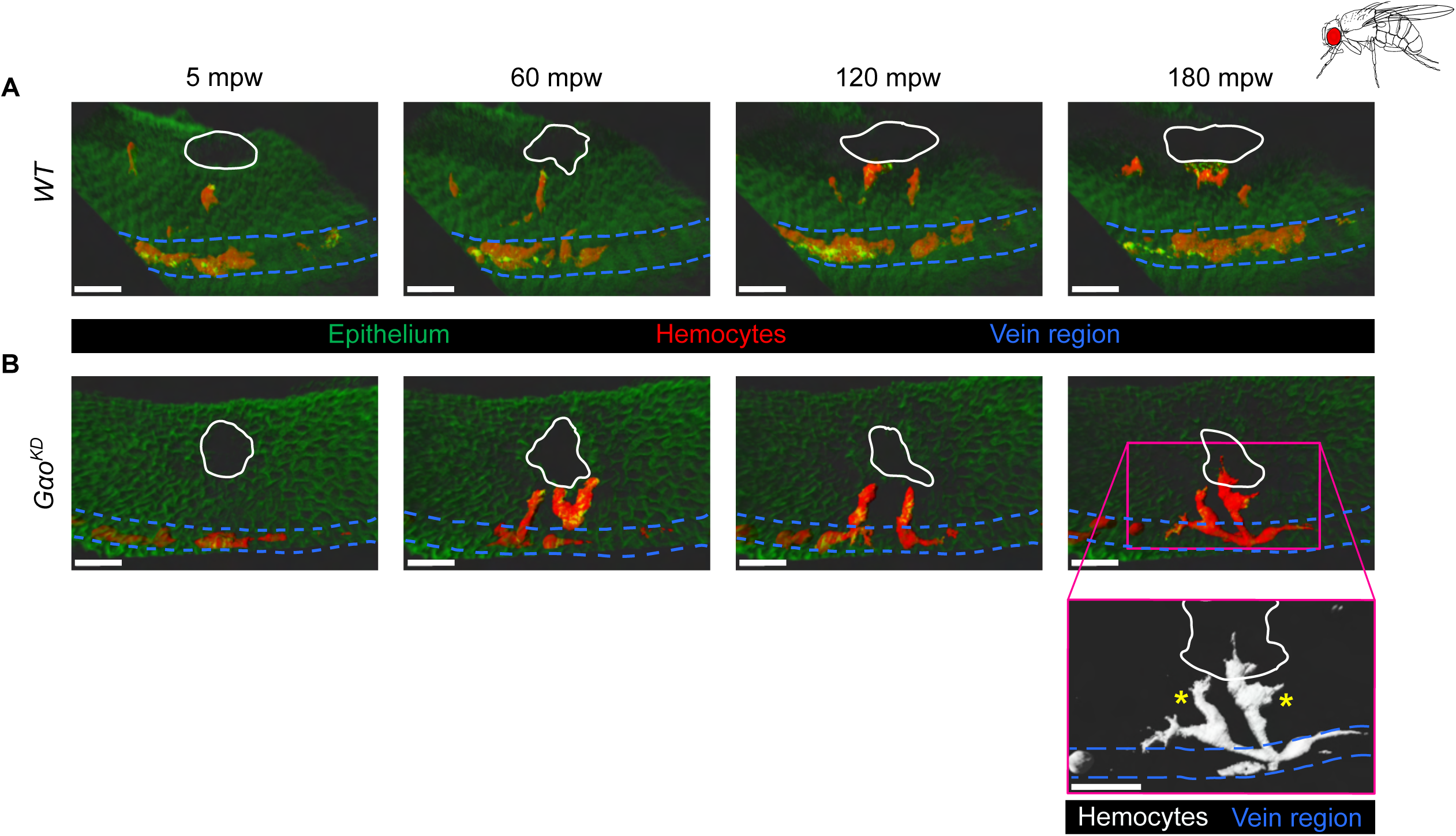
Downstream effectors of Tre1. **(A** and **B)** Representative *in vivo* confocal time-lapse imaging of 75h APF of RNAi-mediated silencing of *Gαo* within hemocytes (*srp-Gal4*-driven knockdown) results in a significant reduction in hemocyte extravasation capacity. Solid white line indicates border of wound, and yellow asterisks indicate extravasating hemocytes. Scale bars: A and B = 15 µm.

## Supplemental Movie Legends

**Supplementary Movie S1. Hemocytes lacking Tre1 are impaired in extravasation.** Related to Figure 1. Representative *in vivo* time-lapse imaging of 75h APF *WT* (left) and *Tre1^KO^* (right) circulating *Drosophila* hemocytes responding and extravasating from pupal veins (blue dotted line) towards sites of laser-induced wounding (solid white line) Examples of extravasating hemocytes are highlighted with white arrows. Wing epithelium is labeled using ubiquitously expressed GMA (green), and hemocytes are labeled by both *srp-Gal4*–driven and the *srp* enhancer trap-driven expression of cytoplasmic mcherry (red). Scale bars represent 20 µm. Z-stacks were taken at regular 4.5 minute time intervals.

**Supplementary Movie S2. Tre1 vs Gpr84 share sequence and structural homology.** Related to Figure 2. Structural comparison of Tre1 (maroon) and its human orthologue GPR84 (teal) with superimposed orthologues highlighting conserved structure. AlphaFold 3 and ChimeraX was used for orthologue structure comparisons.

**Supplementary Movie S3. *Gpr84^-/-^* mice show defective extravasation in TNF-stimulated cremaster muscles.** Related to Figure 3. Representative brightfield intravital microscopy movies of post-capillary venule segments of TNF-stimulated cremaster muscles of WT and *Gpr84^-/-^* mice used to observe leukocyte responses (adhesion and extravasation). Arrows indicate extravasated neutrophils in the interstitial tissue at the 4 h experimental endpoint. Scalebars represent 50 µm.

**Supplementary Movie S4. *Gpr84^-/-^* mice display aberrant extravasation in TNF-stimulated cremaster muscles at the pericyte layer.** Related to Figure 4. Representative confocal intravital microscopy movies of post-capillary venule segments of TNF-stimulated cremaster muscles from WT and *Gpr84^-/-^* on a *Lyz2-EGFP-ki*; *Acta2-RFPcherry-Tg* genetic background; endothelium (blue), pericytes (red) and neutrophils (green). The movie shows a post-capillary venule segment with high optical imaging of a neutrophil breaching the pericyte layer without pause in a WT (taking ∼4 mins) and subsequently a neutrophil exhibiting multiple oscillations at the pericyte layer (*Gpr84^-/-^*) throughout the 1 h imaging period. Scalebars represent 10 µm.

**Supplementary Movie S5**. ***Gpr84^-/-^* mice display impaired detachment from acutely inflamed vessel walls.** Related to Figure 5. Representative confocal intravital microscopy movies of post-capillary venule segments of TNF-stimulated cremaster muscles of WT and *Gpr84^-/-^* on a *Lyz2-EGFP-ki*; *Acta2-RFPcherry-Tg* genetic background; endothelium (blue) and neutrophils (green). Movie shows extravasated neutrophil displaying short uropods prior to detachment in a WT vessel segment. The movie then displays a *Gpr84^-/-^* post-capillary venule segment with neutrophils exhibiting morphologies associated with impaired detachment. Specifically, arrows indicate the formation of elongated uropod structures, persistent uropods and amoeboid neutrophils displaying perivascular arrest over a 1 h imaging period. Scalebars represent 20 µm.

**Supplementary Movie S6. Hemocytes require Gαo for vein extravasation.** Related to Figure 7. Representative *in vivo* time-lapse imaging of 75h APF control (left) and *srp-Gal4* driven *Gαo-RNAi* (right) circulating *Drosophila* hemocytes responding and extravasating from pupal veins (blue dotted line) towards sites of laser-induced wounding (solid white line). Hemocytes expressing *Gαo-RNAi* failed to fully extravasate from vessels over the 3 hours of imaging (right) and remained adhered to vein cells. Wing epithelium is labeled using GFP-tagged E-cadherin (green), and hemocytes are labeled by both *srp-Gal4*–driven and the *srp* enhancer trap expressing cytoplasmic mcherry (red). Scale bar represents 20 µm. Z-stacks were taken at regular 5 min time intervals.

**Supplementary Movie S7. Tre1 is required for Rho1 localization at lagging tails of extravasating hemocytes.** Related to Figure 7. Representative *in vivo* time-lapse imaging of 75h APF control (left) and *Tre1^KO^* (right) *Drosophila* hemocytes extravasating from pupal veins (blue dotted line) towards sites of laser-induced wounding. Upon exit from veins, control hemocytes (left) develop a timely accumulation of active Rho1 that dissipates with de-adhesion to the vessel (white arrow). *Tre1^KO^* hemocytes (right) fail to de-adhere with no observable active Rho1 accumulation at the tail. Active Rho1 is labelled using the Rho1 biosensor (Munjal et al. 2015), and hemocytes are labeled using the *srp* enhancer trap expressing cytoplasmic mcherry (red). Scale bars represent 5 µm. Z-stacks were taken at regular 20 s time intervals.

## Materials and Methods

### *Drosophila* stocks and husbandry

Fly stocks were maintained according to standard protocols. All crosses were performed at 25 °C unless otherwise stated. The following *Drosophila* stocks were used: *ubi-GMA, srp-gal4 (Brückner et al., 2004), UAS-mcherry, srp-mcherry (gift from Daria Siekhaus)*, *Act-Gal4, UAS-GMA, w;;Tre1-RNAi* P{TRiP.HMS00599} (Bloomington *Drosophila* Stock Center #33718), *Oregon R*, *AniRBD* (a gift from T. Lecuit (Munjal et al., 2015)), *w;Tre1^KO^* (a gift from D.J. Andrew (Kim et al., 2021)), *w;;UAS-mys* (Bloomington *Drosophila* Stock Center #68158), *w;Gαo-RNAi* P{TRiP.hms01129} (Bloomington *Drosophila* Stock Center #34653). Of note, in all studies conducted in *Drosophila*, all driver and reporter lines were out-crossed to *Oregon R* (*wild-type*) and the offspring used as a comparative baseline (control) against mutants.

### *Drosophila* dissections, wounding and imaging

Larvae were sexed and male pre-pupae were collected 75 h prior to wounding and kept at 25°C to ensure consistent staging. 75h after puparium formation (APF) pupae were then placed on double sided tape ventral side down and forceps were used to remove the pupa. For imaging, dissected pupae were placed left side down in a glass bottom microscopy dish (Greiner Bio-One GmbH, 627861). Wounds were induced using a Micro Point ablation laser (Andor Technology Oxford Instruments) either attached to an Olympus BX53 upright microscope or a Leica SP8 confocal microscope.

### Identifying Tre1 orthologues, multiple sequence alignment and phylogenetic-tree construction

Tre1 paralogues and orthologues in *Drosophila*, mice and humans were identified using DIOPT with those sequences having a weighted score >5 selected for further analysis (Hu et al., 2011). *Drosophila* protein sequences were retrieved from FlyBase, and human and mouse from NCBI protein database. The longest isoform was selected if there were multiple isoforms per gene. Multiple sequence alignment was performed with Clustal Omega and phylogenetic tree constructed with JalView and Interactive Tree of Life (iTOL). Alignment and visualization of orthologous proteins structures was performed using UCSF ChimeraX (Pettersen et al., 2021).

### RNA isolation, reverse transcription, and real-time quantitative PCR

RNA was isolated from *WT* and *Tre1^KO^;;* 75 h APF pupae by crushing in TRIzol (Life Technologies) and RNA purified using an RNeasy Mini Kit (Qiagen). Equal quantities of RNA were then reverse transcribed using a Thermo Scientific Maxima First Strand cDNA Synthesis Kit, and genomic DNA was eliminated using double strand-specific DNase (Thermo Scientific). Relative quantification of gene expression was performed using SYBR Green Supermix with a real-time PCR machine. Target gene expression was normalized to the expression of the housekeeping reference gene *Gapdh1* using the ΔΔCt analysis method. The following primers were used in this study: *Tre1* F-primer 5′-ATTAGTGCCTGTGTCTTTGTGAC-3′ and R-primer 5′-GGAGATGCTTAGCGAAATGACG-3′, *mys* F-primer 5′-CGCGGTGCTACCAAAACAC-3′ and R-primer 5′-GAATCTGCTCAACTGTTATCGGA-3′, *Fak* F-primer 5′-GCTGACCGATGATTATGC-3′ and R-primer 5′-CGAACGGTGGGCGTAGAGTAG-3′, and *Gapdh1* F-primer 5′-TAAATTCGACTCGACTCACGGT-3′ and R-primer 5′-CTCCACCACATACTCGGCTC-3′.

### Correlative Light Electron Microscopy on pupae

For CLEM studies, confocal images were acquired as described and wing hinges were punctured with forceps before fixation in freshly prepared 2.5% glutaraldehyde (Thermo Fisher Scientific) in 0.1M cacodylate buffer (Thermo Fisher Scientific) for 24 hours at 4°C. Pupae were then washed in 0.1 M sodium cacodylate (three times for 10 min) before being placed in a secondary fixative containing 1% OsO_4_, 1.5% potassium ferrocyanide and 0.1 M sodium cacodylate for 2 h at 4°C. They were again washed in 0.1 M sodium cacodylate (three times for 10 min) and wings dissected and embedded in a solution of 13.5% BSA, 0.1 M cacodylate and 5% glutaraldehyde for 10 minutes. Wings were then placed in an automatic tissue processor (Leica EM TP). 70 nm sections were cut with a UC6 ultramicrotome (Leica) with Diamond Knife (Diatome), and were collected onto pioloform-coated copper slot grids (Agar Scientific). These sections were then imaged using a Tecnai 12 BioTwin Spirit TEM (tungsten filament, 120 kV) and a FEI Eagle 4k×4k CCD camera.

### Animals

All transgenic mice used were on a C57BL/6N background. *Gpr84^-/-^* mice (and WT littermate controls) were derived from cryopreserved sperm (provided by GlaxoSmithKline) at MRC Harwell. GPR84 (GenBank accession number NM_030720, Ensembl identification number ENSMUSG00000063234) KO mice were provided by Deltagen under a GSK license agreement. As summarised in (Nicol et al., 2015), the insertion of a an IRES–LacZ–poly(A) expression cassette replaced 257 bp of coding sequence to generate a premature stop codon, thus deleting in the first three transmembrane domains. Heterozygote breeding pairs were used to maintain the colony at Queen Mary University of London.

For confocal intravital microscopy studies, myeloid cells and pericytes were visualised by generating a *Lyz2-EGFP-ki*; *Acta2-RFPcherry-Tg* colony. The Lyz2-EGFP-ki mice, summarised in (Faust et al., 2000), were crossed with *Acta2-RFPcherry-Tg* mice in which insertion of the RFP variant mCherry under control of the *Acta2* promotor to generate RFP+ pericytes and smooth muscle cells (Proebstl et al., 2012). *Lyz2-EGFP-ki*; *Acta2-RFPcherry-Tg* mice were crossed with *Gpr84^+/-^* (heterozygote) mice across several generations to generate WT and *Gpr84^-/-^* mice on the *Lyz2-EGFP-ki*; *Acta2-RFPcherry-Tg* background.

For the chimeric studies detailed below, *Lyz2-EGFP-ki* mice were used as bone-marrow donors to C57Bl6/J recipient mice with a CD45.1 allelic variant, which were a gift from Dr Suchita Nadkarni (William Harvey Institute, Queen Mary University London).

All animals were used between 8-20 weeks old, and all experimental groups were age matched. For all pharmacological blockade studies, wildtype C57/Bl6J mice were purchased from Charles River. Male mice were used for all cremaster and peritonitis studies; both males and females were used for ear IR and cutaneous wound healing experiments.

All animals were group housed in individually ventilated cages under specific pathogen free (SPF) conditions and a 12 h light-dark cycle. Animals were humanely sacrificed via cervical dislocation or exsanguination at the end of experiments in accordance with UK Home Office regulations. All *in vivo* experiments were conducted at the William Harvey Research Institute, Queen Mary University of London, UK under the UK legislation for animal experimentation and in agreement with the UK Home Office Animals Scientific Procedure Act 1986 (ASPA).

### Brightfield Intravital microscopy and analysis

Cremaster muscles were inflamed with an intrascrotal (i.s.) injection of TNF (300 ng) or vehicle control in a 400 µl bolus of PBS. The mice were administered an i.p. injection of ketamine (100 mg/kg) and xylazine (10 mg/kg) 1.5 h prior to exteriorisation of the cremaster muscle onto an imaging stage. The cremaster muscles were continuously superfused in warm Tyrode’s solution and the animals were maintained at 37 °C throughout image acquisition.

Brightfield intravital microscopy was used as a preliminary assessment of real-time leukocyte dynamics. Firmly adhered leukocytes were considered those that remained stationary on the luminal endothelium for ≥ 30 s. The number of extravasated leukocytes within 50 µm of a 100 µm post-capillary venule segment was quantified at the 4 h experimental end point. Post-capillary venules were observed and movies captured using a 63x water dipping objective on a transmitted light upright fixed stage microscope (Axioskop FS,Carl Zeiss) with a digital CMOS camera (Hamamatsu).

### Generation of bone marrow chimeric animals

To generate chimeric mice, C57Bl6/J mice (on a CD45.1 allelic background) were irradiated with 4 Gy (3 min 25 s) two times, 4 hours apart. The day after irradiation, ∼1.5 x 10^6^ bone marrow derived cells were administered to each mouse by i.v. injection. 4 weeks after irradiation, myeloid cell reconstitution efficiency was assessed by whole blood collection and flow cytometry to calculate absolute leukocyte counts prior to induction of inflammation in the cremaster muscles (all mice showed an effective reconstitution efficiency shown in **Supplementary Figure 2**). After 4 h the animals were sacrificed, the cremaster muscles were exteriorised and fixed prior to permeabilization and immunofluorescence staining.

### Inflammatory response in cremaster muscles

Mice were briefly anaesthetised with 3% isofluorane and injected i.s. with TNF (300 ng) (R&D systems) or phosphate buffered saline (PBS) as a vehicle control, together with fluorescently labelled non-blocking anti-CD31 mAb (4 µg, clone 390, Thermo Fischer Scientific) to label endothelial cell junctions, in a 400 µl bolus using a 2 – 5 h stimulation periods.

### Whole mount immunofluorescence staining

Exteriorised cremaster or ear tissues were fixed for 1 h in 4 % paraformaldehyde (PFA) washed in PBS and permeabilised for 3 or 5 h, respectively, in PBS containing 25 % fetal calf serum (FCS) and 0.5% Triton-X. Tissues were stained with fluorescently labelled monoclonal antibodies, MRP-14 (a gift from Professor Nancy Hogg) or PECAM-1, overnight in PBS containing 10 % FCS at 4 °C. Antibody conjugation to AF488, 647 or 405 was performed using kits according to the manufacturer’s instructions (ThermoFisher Scientific).

### Confocal microscopy and image analysis

Immunostained whole-mount cremaster muscles or ears were imaged with an up-right Leica TCS SP8 (Leica) (with a white light laser (tuneable 470–670 nm)) or inverted LSM800 (Carl Zeiss) confocal laser scanning microscope equipped with solid-state laser diodes (405, 488, 561 and 640 nm excitation wavelengths), respectively. For high resolution confocal imaging of neutrophil-vessel wall structures, either the LSM800 (Carl Zeiss) or a CSU-W1 SoRa Spinning Disk microscope (Nikon) was used prior to deconvolution with either Huygens 25.10 (Scientific Volume Imaging) or NIS version 6.10.02 (Nikon), respectively.

For all quantified images, Z-stacks of half post-capillary venules, in a longitudinal orientation, were collected. Post-capillary venules 20-45 µm diameter were selected for image acquisition and images collected at 1024 x 1024 pixels. In cremaster tissues, the number of extravasated neutrophils were quantified per field of view (typically 330 x 160 x 45 µm). For IR injury experiments in the ear (described below), the number of extravasated neutrophils within 50 µm from the vessel wall was quantified.

### Confocal intravital microscopy and analysis

Confocal IVM was used to assess the migration dynamics of neutrophils across the endothelium and interactions with pericytes. WT and *Gpr84^-/-^* mice on a *Lyz2-EGFP-ki*; *Acta2-RFPcherry-Tg* mice, mice were subject to an injection of TNF (300 ng). Movies were acquired on a Leica SP8 laser scanning microscope with a 20x water-dripping objective (NA 1.0) and 8 kHz resonant scanner. Postcapillary venules between 20-45 µm were imaged for 1 h with two venules imaged per mouse 2-5 h post TNF-injection. Half vessels in a longitudinal orientation were recorded to achieve dimensions of typically 300 x 130 x 35 µm with a resulting voxel size of approximately 0.29 (x) x 0.29 (y) x 0.69 (z) µm and serial z-stacks of 0.7 µm optical sections were acquired every 60 s. The time-lapse serial z stacks were assembled and processed into 3D videos and rendered using IMARIS software^TM^ (Bitplane, Zurich, Switzerland) for analysis.

Neutrophil dynamics were determined by manual tracking of individual neutrophils using IMARIS software. Neutrophil TEM was defined as an event where neutrophils fully migrated through EC junctions in a luminal-to-abluminal direction. Trans- pericyte migration (TPM) was defined as the point at which a neutrophil displayed an initial protrusion through the pericyte layer before completing their migration into the perivascular space. Neutrophils were categorised as displaying “incomplete” TPM when they displayed a minimum of 1 oscillation during the 1 h imaging period - a protrusion of the leading edge through the pericyte border followed by a retraction into the sub-endothelial space. In **Figure 4B**, the maximum length of a peri-vascular neutrophil was measured within a 1 h imaging period; measurements were taken from the leading edge of the neutrophil to the point of attachment between the posterior protrusion and the venular wall. In **Figure 4C**, the time attached to the vessel wall was considered the period between the end of TPM and the completion of tail detachment from the venular wall.

### Transmission Electron Microscopy

For mouse transmission electron microscopy studies, cremaster muscles were excised from mice subjected to 4 h TNF induced inflammation, pinned to dental wax before fixation in a freshly prepared mix of 2.5% glutaraldehyde (Thermo Fisher Scientific) in 0.05 M cacodylate buffer (Thermo Fisher Scientific) supplemented with 0.18 mM calcium chloride, for 16 hours at 4°C. Samples were then secondarily fixed in 1% OsO_4_ for 1 hour, and taken through serial dehydrations in an automatic tissue processor (Leica EM TP) prior to embedding in resin. 70 nm sections were cut with a UC6 Leica ultramicrotome and Diamond Knife (Diatome) and were collected onto pioloform-coated copper slot grids (Agar Scientific) and imaged using a Tecnai 12 BioTwin Spirit TEM (tungsten filament, 120 kV) and a FEI Eagle 4k×4k CCD camera.

### Whole blood collection

Mice were either terminally anesthetized with 3% isoflurane and the abdomen exposed to collect whole blood from the caudal vena cava followed by cervical dislocation or sacrificed prior to a cardiac puncture with collection of whole blood directly into 50 mM EDTA.

### Flow cytometry

Differential leukocyte counts in whole blood and peritoneal lavage were assessed by flow cytometry. Blood samples were incubated with ACK erythrocyte lysis buffer (150 mM NH_3_CL, 1 mM KHCO_3_ and 1 mM EDTA) for 5 min. Samples were then washed once with staining buffer (2 mM EDTA and 0.5% BSA in PBS). Samples were then incubated with Fc-block (anti-CD16/32) antibodies (5 µg/ ml) diluted in staining buffer for 15 min at 4 °C. Subsequently samples were stained for surface cells markers diluted in staining buffer (and their appropriate isotypes controls); anti-CD45 PB (1 µg/ ml; 30-F11; 103126; BioLegend), CD3 PE (1 µg/ ml; 17A2; 100205; BioLegend), CD19 AF488 (1 µg/ ml; 6D5; 115524; BioLegend), CD11b PE/Cy7 (0.2 µg/ ml; M1/70; 101216; BioLegend), Ly6C PerCP-Cy5.5 (0.2 µg/ ml; HK1.4; 128012; BioLegend), Ly6G AF647 (0.5 µg/ ml; 1A8; 127610; BioLegend), F4/80 PE (1 µg/ ml; W20065B; 123117; BioLegend), CD115 APC-Cy7 (1 µg/ ml; AFS98; 135531; BioLegend), CD62L PE (0.2 µg/ ml; MEL-14;104407; BioLegend), CD11a PE (0.1 µg/ ml; M17/4; 101107; BioLegend), CD11b PE (0.1 µg/ ml; M17/0; 101207; BioLegend), CD61 PE (0.4 µg/ ml; 2C9.G2 (HMβ3-1); 101107; BioLegend), CD51 PE (0.4 µg/ ml; RMV-7; 104105; BioLegend), CD29 PE (0.4 µg/ ml; HMβ1-1; 102207; BioLegend), CD18 PE (2 µg/ ml; M18/2; 101407; BioLegend), CD49 PE (0.4 µg/ ml; 5H10-27 (MFR5); 103805; BioLegend), CD11c PE (1 µg/ ml; N418; 117307; BioLegend). Gating strategies were based on isotype and FMO controls. Accurate cell counts were validated using counting beads and analysed using a LSR Fortessa flow cytometer (BD Biosciences) and FloJo software (TreeStar).

### Bulk-RNA sequencing

Neutrophils from the whole blood of male WT versus GPR84 KO mice were FACs sorted and gated using Zombie Yellow™ fixable Viability kit as a live/dead marker, single cells, CD45^+^, CD3^-^, CD19^-^, CD11b^+^, Ly6C^int^, Ly6G^+^ using a FACS Aria™ II (BD Biosciences). Exactly 10,000 neutrophils were collected in Trizol and sent to GENEWIZ (Azenta Life Sciences, US) for RNA isolation, quality control and ultra-low input RNA sequencing amplification. Sequencing was performed on an Illumina HiSeq instrument with a 2x150-base-pair (BP) paired end configuration. 35 million reads were mapped to the reference mouse transcriptome (Gencode release v28, GRCm39) and transcripts were quantified by STAR aligner v2.7.3a in 2-pass mapping mode. Protein coding and immunoglobulin genes were used for analysis. Counts were subjected to variance-stabilising transformation (VST) using DESeq2 v.1.42.1. Principal component analysis (PCA) was undertaken with prcomp function in R using the VST data and its plots were generated using ggplot2 v3.4.4. Differential expression analysis was performed on count data using DESeq2 v.1..42 using an unadjusted p-value cut-off of 0.05 to identify transcripts differentially expressed between genotypes, where the volcano plot was visualized using easylabel v0.3.3. Pathway analysis was performed on DE transcripts using the WikiPathways 2024 by EnrichR database (as described by (Kuleshov et al., 2016) with enriched pathways considered significant with an adjusted p-value of P < 0.05. Heatmaps were generated using ComplexHeatmap v2.21.1.

### *In vitro* adhesion assay

Ibidi chamber 8-well slides were coated overnight with 3 ug/ml rmICAM-1 (Abcam) at 4 °C, blocked with 1% Casein (in PBS) for 1 h at room temperature (RT) and washed with PBS. Murine neutrophils were isolated from bone marrow cells (harvested from tibiae and femora) by negative magnetic cell sorting using the Neutrophil isolation kit (Miltenyi Biotec) according to the manufacturer’s instructions. The purity of isolated neutrophils (CD45^+^/Ly6G^+^) expression profile) was consistently > 95% as determined by flow cytometry. Isolated neutrophils were stained with Cell Tracker™ Green CMFDA or Cell Tracker™ Orange CMRA (Thermofisher Scientific) at 0.5 µM for 10 mins at RT. Potential effects of Cell Tracker™ colours were controlled for by alternating between genotypes for experimental replicates. Isolated neutrophils were seeded at 100,000 of each genotype and left to settle for ∼ 20 min at RT prior to stimulation with 10 nM CXCL1 (Peprotech) for 15 mins at 37 °C and imaged on a Nikon Spinning Disk SoRa at 60x magnification prior to assessment of the proportion of “spread” or motile cells by manual counting on IMARIS software. Neutrophils were classified as “spread” when displaying a flattened and non-amoeboid like structure and considered as “motile” when displaying membrane ruffling and migration on the ICAM-1 coated surface.

### Zymosan-induced peritonitis

C57Bl6/J mice (Charles River) received 10 or 30 mg/ kg GLPG1205 (Galapagos Pharmaceuticals) in (40% PEG 200, 60% sterile water) or vehicle control by oral gavage 24 h prior to induction of inflammation in the TNF-inflamed cremaster model or zymosan-induced peritonitis. Peritonitis was induced by an i.p. injection of zymosan (1 mg) in 500 µl of PBS or PBS alone as control. Subsequently, 24 h later, the peritoneal lavage was collected in 5 ml PBS containing 5mM EDTA (Sigma-Aldrich) and 0.25% BSA. The peritoneal exudates were labelled for myeloid cell markers of interest for flow cytometry and gated as follows for neutrophils (single cells, live, CD45^+^ CD115^-^ F4/80^-^ Ly6C^int^ Ly6G^+^) and monocytes (single cells, live, CD45^+^ CD115^+^ F4/80^low-int^ Ly6C^hi-int^ Ly6G^-^). Absolute cell counts were calculated with the application of counting beads and assessed by flow cytometry.

### Ischaemia- reperfusion injury in the ear

Male and female C57Bl6/J mice (Charles River) received 30 mg/ kg GLPG1205 or vehicle control as above. Mice were briefly anaesthetised with 3% isoflurane prior to application of magnets covering the lower region of the ear; anaesthesia was reduced to 1.5 % during the 1.5 h ischaemia period and a heating pad used to maintain body temperature at 37 °C. The magnets (2mm thick 45H Neodymium with 0.8 kg pull) were removed prior to a 3 h reperfusion period. Mice were sacrificed, whole blood collected (to assess differential leukocyte counts) and ear tissues harvested and processed for staining and analysis.

### Cutaneous wound injury

Male and female C57Bl6/J mice (Charles River) received 30 mg/ kg GLPG1205 or vehicle control as above. Briefly, 8-week-old mice were anaesthetised with 3% isofluorane, prior to shaving their back skin and generating paired excisional wounds with a 4 mm biopsy punch (Kai Medical). Animals were sacrificed at 0, 18 and 36 h post-wounding, and skin samples were processed for histological analysis.

### Histological staining and image analysis

Tissue was fixed in 4% paraformaldehyde overnight prior to transfer through serial dehydration steps in preparation for embedding in paraffin using an automatic tissue processor (Expredia Exelsior AS). 7 μm sections were cut with a Leica 2125 microtome and subsequently deparaffinised and stained with Haematoxylin and Eosin. Stained sections were then imaged in an Olympus VS200 slide scanner, followed by analysis in QuPath (Bankhead et al., 2017). Quantification of the total number of neutrophils was performed using the built-in Cell Detection algorithm that detects haematoxylin-stained nuclei with minor adjustments to intensity parameters for neutrophil nuclear morphology: threshold (0.15) and cell expansion (2 µm). This was validated by comparing manual and automated counts.

### Quantification and statistics

Statistical analyses were performed using Prism 10.1 software (GraphPad) and all datasets are expressed as mean ± SEM. Details for exact n numbers are reported in the legends for each Figure. Sample sizes were chosen based on published work in which similar results have been reported.

## Acknowledgements

We would like to thank members of the Weavers, Martin and Nourshargh labs for helpful discussion. We would like to thank the technical assistance of Laura Vazquez Martinez, Navid Mousavi and Mohammed Yaseen. We also thank the Wolfson Bioimaging Facility (University of Bristol, UK), Bloomington Stock Centre (University of Indiana, USA) as well as the BCI Flow Cytometry Facility and BSU Animal Facility at Queen Mary University of London, UK. This research was funded by an MRC Programme Grant to P.M., S.N. and H.W. [MR/V011294/1], and Wellcome Trust grants [221699/Z/20/Z] (SN), [217169/Z/19/Z] (PM) and [208762/Z/17/Z] (HW). For the purpose of Open Access, the author has applied a CC BY public copyright license to any Author Accepted Manuscript arising from this submission.

## Author contributions

**Clare Latta^1*^**

Formal analysis, Investigation, Writing - original draft, Writing - review & editing

**Terrence M. Trinca^2*^**

Formal analysis, Investigation, Writing - original draft, Writing - review & editing

**Francesca Robertson^2^**

Investigation

**Rachel Lau^1^**

Formal analysis

**Anna Barkaway^1^**

Investigation

**Loic Rolas^1^**

Investigation

**Matthew Golding^1^**

Investigation

**Paul Imbert^1^**

Visualisation, Resources

**Barbara Walzog^4^**

Investigation

**Almke Bader^4^**

Investigation

**Liam S. Hill^2^**

Investigation

**Lorna Hodgson^2^**

Investigation

**Myles Lewis^1^**

Resources

**Mathieu Benoit-Voisin^1^,**

Methodology, Resources, Supervision, Visualization, Writing - original draft, Writing - review & editing

**Paul Martin^2^,**

Conceptualization, Funding acquisition, Methodology, Project administration, Resources, Supervision, Visualization, Writing - original draft, Writing - review & editing

**Sussan Nourshargh^1^**

Conceptualization, Funding acquisition, Methodology, Project administration, Resources, Supervision, Visualization, Writing - original draft, Writing - review & editing

**Helen Weavers^2^**

Conceptualization, Funding acquisition, Methodology, Project administration, Resources, Supervision, Visualization, Writing - original draft, Writing - review & editing

## Data availability statement

RNA-Seq data is available at ArrayExpress (with accession ID E-MTAB-16278) and the remaining manuscript data will be made available upon reasonable request to the senior authors.

